# Cytosolic sodium accumulation is a common danger signal triggering endocytic dysfunction and NLRP3 inflammasome activation

**DOI:** 10.1101/2024.03.26.586774

**Authors:** Matthew S.J. Mangan, Anil Akbal, Sophie Rivara, Agnes Schröder, Gabor Horvath, Philipp Walch, Jan Hegermann, Romina Kaiser, Fraser Duthie, Joel Selkrig, Jonathan Jantsch, Ralf Gerhard, Dagmar Wachten, Eicke Latz

**Affiliations:** Institute of Innate Immunity, University Hospital Bonn, University of Bonn, 53127 Bonn, Germany; German Center for Neurodegenerative Diseases, 53127, Bonn, Germany; Department of Orthodontics, University Hospital Regensburg, 93053, Regensburg, Germany; Institute for Medical Microbiology and Hygiene, University Hospital Regensburg, 93053, Regensburg, Germany; European Molecular Biology Laboratory (EMBL), Genome Biology Unit, 69117 Heidelberg, Germany; Heidelberg University, Faculty of Biochemistry, 69120 Heidelberg, Germany; Institute of Functional and Applied Anatomy, Research Core Unit Electron Microscopy, OE8840, Hannover Medical School, 30625, Hannover, Germany; Institute for Medical Microbiology, RWTH University Hospital, 52074 Aachen, Germany; Institute for Medical Microbiology, Immunology and Hygiene, Center for Molecular Medicine Cologne, University Hospital Cologne and Faculty of Medicine, University of Cologne, 50935, Cologne, Germany; Institute of Toxicology, Hannover Medical School, 30625 Hannover, Germany; Deutsches Rheuma Forschungszentrum Berlin (DRFZ), 10117, Berlin, Germany; Department of Infectious Diseases & Immunology, UMass Medical School, Worcester, MA 01605, USA

**Keywords:** NLRP3, inflammasome, sodium, potassium, Toxin B, TcdB, Clostridioides difficile, lysosome, cell volume

## Abstract

Detecting and responding to noxious molecules internalized within the endolysosomal system, including bacterial toxins and particulate matter, is essential to prevent cellular intoxication and damage. Here, we demonstrate that the NLRP3 inflammasome detects perturbations of the endolysosomal system by large clostridial toxins, including toxin B from *Clostridioides difficile*, as well as monosodium urate and silica crystals in human macrophages. These molecules cause sodium efflux from the endolysosomal system into the cytosol, driving cytosolic sodium accumulation. The rapid increase in cytosolic sodium subsequently triggers cell swelling and inhibits endocytic trafficking to activate the NLRP3 inflammasome. Furthermore, we demonstrate that cytosolic sodium accumulation is a common trigger for NLRP3 activation by non-particulate stimuli, including nigericin and inhibition of the Na^+^/K^+^ ATPase. Our findings reveal that accumulation of cytosolic sodium is the common denominator underlying activation of the NLRP3 inflammasome upon exposure to different danger signals.

## Introduction

The endolysosomal system serves as an entry route for noxious molecules, including bacterial toxins, pathogens, and particulate matter, many of which perturb endocytic membranes to enter the cytoplasm ^1–3^. However, how these perturbations are detected by the cell remains poorly understood. Particulate matter, including crystals and protein aggregates, can damage and rupture lysosomal membranes, leading to cathepsin release into the cytosol ^1^. However, most pathogen-derived molecules translocate across endocytic membranes without causing gross membrane damage ^2^, and it remains unclear whether such events are sensed through alternative mechanism(s). An important group of disease-causing toxins which gain entry to the cytoplasm through the endosomal system are the large clostridial toxins (LCT) ^4^. These include toxin B (TcdB) and toxin A (TcdA) from *Clostridioides difficile*, which causes colitis and even sepsis solely through expression of these toxins ^5^.

In murine macrophages, TcdB triggers Pyrin inflammasome activation by inhibiting RhoA/B/C ^6^. In contrast to this, we discovered that TcdB is sensed by the NLRP3 inflammasome in human monocyte-derived macrophages (hMDM), as the Pyrin inflammasome activation requires additional priming steps in hMDM ^7^. However, the underlying molecular mechanisms remain ill-defined. NLRP3 inflammasome activation is a two-step process: It first requires a “priming” step, whereby activation of toll-like and cytokine receptors induce NLRP3 expression ^8^ and changes in post-translational modification ^9^, and a second ‘activation’ step, which then triggers inflammasome assembly ^10^. A decrease in the intracellular potassium concentration is considered essential for most NLRP3 activators ^11^, though it remains unknown how this facilitates NLRP3 activation. Additionally, NLRP3 activation is thought to require disruption of the trans-Golgi network (TGN) or endocytic trafficking ^12,13^, though it is unclear how physiologically relevant NLRP3 stimuli would trigger dysfunction of either system, as these studies were mostly limited to activation of NLRP3 by small molecules such as R837 and ionophores, including nigericin ^12,13^.

To gain insights into the activation mechanism of NLRP3, we deciphered how TcdB activates the NLRP3 inflammasome. We determine that TcdB, as well as monosodium urate (MSU) and silica crystals, trigger NLRP3 inflammasome activation by promoting sodium translocation from the endolysosomal network into the cytosol causing cytosolic sodium accumulation. This rapid osmotic shift induced cell swelling and disrupted the endocytic network, leading to activates NLRP3. Furthermore, we establish that cytosolic sodium accumulation is a common requirement for NLRP3 inflammasome formation by diverse NLRP3 activators, including the ionophore nigericin and the Na^+^/K^+^ ATPase ouabain. Markedly, cytosolic sodium accumulation occurs subsequently to or independently from cytosolic potassium efflux, which was previously considered to be required for NLRP3 activation. Thus, we establish that rapid cytosolic sodium accumulation is a common denominator for NLRP3 activation.

## Results

### Large clostridial toxins activate NLRP3 by structural perturbation

During intoxication, TcdB is internalized by receptor-mediated endocytosis, then undergoes a confirmational change at acidic pH (∼5.2), inserts into the endolysosomal membrane, and translocates to the cytosol ^14^. The glucosyltransferase domain (GTD) of the toxin is released in the cytosol, where it glucosylates and, in turn, inactivates RhoA/B/C, Rac1/2/3, and Cdc42 ^14^ (Fig. 1A).

**Figure 1.**
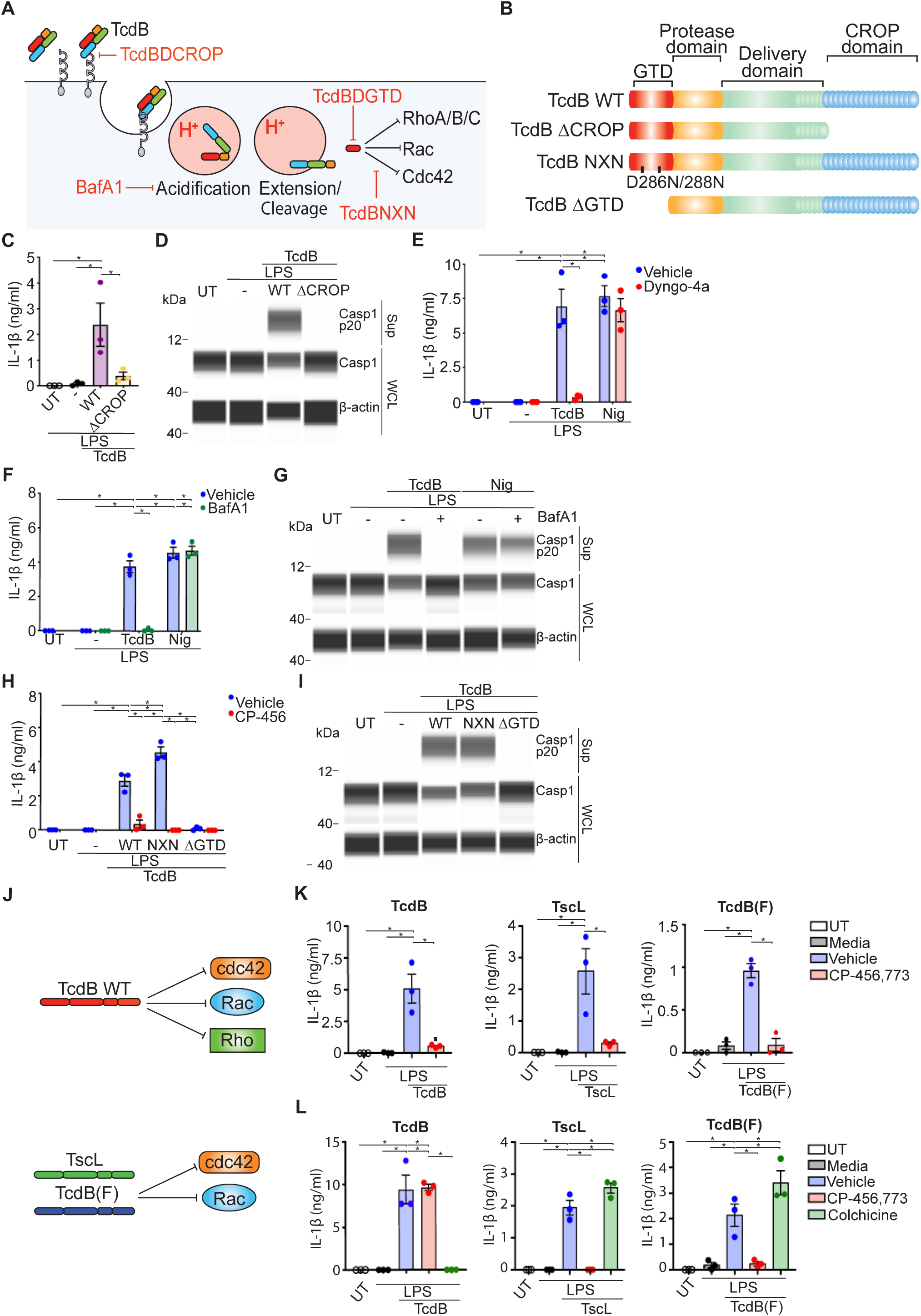
Activation of NLRP3 by TcdB requires receptor-mediated uptake and endosomal acidification. **A)** TcdB cellular entry and intoxication. **B)** TcdB variants used in this study. LPS-primed hMDM (10 ng/ml, 3 h) incubated with TcdB or TcdB ΔCROP (20 ng/ml, 2h). IL-1β was assessed from cell-free supernatants (sup) **(C)** or caspase-1 was assessed from sup or whole cell lysate (WCL) by simple western assay (SWA) **(D)**. **E)** LPS-primed hMDM with TcdB (20 ng/ml) or nigericin (8 μM) after pre-incubation with Dyngo-4a (20 μM, 30 min). IL-1β was assessed from supernatants. LPS-primed hMDM incubated with TcdB (20 ng/ml, 2 h) or nigericin (8 μM, 2 h) after BafA1 pre-treatment (200 nM, 15 min). IL-1β was assessed from sup **(F)** or caspase-1 was assessed from sup or WCL by SWA **(G)**. LPS-primed hMDM (10 ng/ml, 3 h) incubated with TcdB, TcdB NXN, or TcdB ΔGTD (20 ng/ml, 2 h) with or without CP-456,773 pre-incubation (2.5 μM, 15 min). IL-1β was assessed from sup **(H)** or caspase-1 was assessed from sup or WCL by SWA **(I)**. **J)** Substrate specificity of TcdB or TcdB(F) and TscL. IL-1β assessed from sup from **K)** LPS-primed hMDM (10 ng/ml, 3 h) or **L)** monocytes (1 ng/ml, 3 h) incubated with TcdB, TcdB(F) (100 ng/ml) or TscL (100 ng/ml) following incubation with CP-456,773 (2.5 μM, 15 min) or colchicine (2 μM, 15 min). All data are shown as mean ± SEM, n = 3, with each point representing one donor. SWAs representative of n = 2. Statistical analysis performed using one-way ANOVA (C, K-L) or two-way ANOVA (E-F, H) *p<0.05, only significant differences annotated.

We hypothesised that TcdB-mediated NLRP3 activation requires receptor binding and subsequent internalization into the endocytic network by clathrinmediated endocytosis ^15,16^. To determine whether receptor binding was required, we used a TcdB mutant lacking the C-terminal combined-repetitive oligopeptides (CROP) domain required for receptor binding (Fig. 1B, S1A) ^16^. In contrast to TcdB WT, the TcdB ΔCROP mutant did not trigger IL-1β release (Fig. 1C), caspase-1 cleavage (Fig. 1D), or IL-1β cleavage (Fig. S1B), all measures of inflammasome activation, demonstrating that receptor binding is required for NLRP3 activation. To confirm that TcdB ΔCROP was not internalized, we assessed glucosylation of Rac2/3 by TcdB using an anti-Rac1/2/3 antibody that fails to recognize its epitope after glucosylation ^17^. As the GTD can only modify Rac2/3 if it reaches the cytosol, assessing Rac2/3 modification acts as a surrogate to determine if TcdB is internalized. Compared to TcdB WT, TcdB ΔCROP did not modify Rac2/3, demonstrating that the GTD does not reach the cytosol (Fig. S1C). We then tested whether subsequent internalization of TcdB was required for NLRP3 activation by inhibiting clathrin-mediated endocytosis with Dyngo-4a, a small molecule dynamin inhibitor. Pre-incubation with Dyngo-4a prevented almost all TcdB uptake (Fig. S1D) and TcdB-mediated IL-1β release (Fig. 1E), whereas NLRP3 activation by nigericin, an ionophore that mediates K^+^/H^+^ exchange, was unaffected (Fig. 1E).

To investigate whether the acidification-mediated conformational change of TcdB in the endo/lysosome, which promotes insertion into the endolysosomal membrane and delivers the GTD to the cytosol ^18^, is required for NLRP3 activation, we applied the V-ATPase inhibitor Bafilomycin A1 (BafA1), which inhibits endolysosomal acidification. BafA1 pre-incubation prevented both TcdB-mediated IL-1β release (Fig. 1F) and caspase-1 cleavage (Fig. 1G), as well as inhibiting glucosylation of Rac2/3 by TcdB (Fig. S1E), demonstrating that it blocked delivery of the GTD to the cytosol. In contrast, BafA1 treatment did not affect NLRP3 activation by nigericin (Fig. 1F-G). These data collectively demonstrate that endosomal acidification is required for TcdB-mediated NLRP3 activation.

The GTD is the effector domain of TcdB, containing the enzymatic motif required for glucosylation and inactivation of RhoA/B/C, Rac1/2/3, and Cdc42^4^. To determine its role in NLRP3 activation, we used two TcdB mutants. The first, TcdB NXN, contained mutations at residues 286/288 that ablated glucosyltransferase activity ^19^. The second mutant, TcdB ΔGTD, lacked the entire GTD (Fig. 1B, S1F). We confirmed that neither mutant glucosylated Rac2/3, demonstrating that they lack glucosyltransferase activity (Fig. S1G). Compared to TcdB WT, loss of GTD enzymatic activity in the TcdB NXN mutant promoted IL-1β release, (Fig. 1H), but did not alter caspase-1 cleavage (Fig. 1I), while the TcdB ΔGTD mutant did not trigger IL-1β release (Fig. 1H) or caspase-1 cleavage (Fig. 1I). The NLRP3 inhibitor CP-456,773 (also known as MCC950 or CRID3) ^20^ inhibited IL-1β release by the TcdB NXN mutant as well as TcdB WT (Fig. 1H), confirming that IL-1β release depended on NLRP3. These results demonstrated that TcdB-mediated NLRP3 activation required the GTD but not its enzymatic activity.

We hypothesized that the insertion of the GTD, which governs the translocation into the cytoplasm, is the structural perturbation that is required to activate NLRP3. As the delivery domain is highly conserved in other LCTs (Schirmer et al. 2004), we hypothesized that NLRP3 is a general sensor of internalized LCT. To test this, we assessed whether TcdB(F), another pathogenic variant of TcdB, previously identified from clinically relevant isolates of *C. difficile,* or Lethal Toxin from *Paeniclostridium sordellii* (TscL) also activate NLRP3. Both toxins inactivate Rac2/3 and Cdc42, but neither inactivate RhoA/B/C (Fig. 1J) ^17^. Similar to TcdB, TcdB(F) and TscL triggered NLRP3-dependent IL-1β release in a CP-456,7773-dependent manner (Fig. 1K), demonstrating that they activate the NLRP3 inflammasome. We then assessed whether LCT triggered an inflammasome response in monocytes. Unlike hMDM, monocytes have an active Pyrin inflammasome ^7^, which is triggered by inactivation of RhoA/B/C ^6^. Based on the substrate specificity, only TcdB should trigger a Pyrin response. Indeed, TcdB activated the Pyrin inflammasome, as IL-1β release was blocked by the Pyrin-inhibitor colchicine (Fig. 1L). In contrast, TcdB(F) and TscL triggered IL-1β release in a NLRP3-dependent manner (Fig. 1L). This indicates that detection of LCT by NLRP3 through structural perturbation insertion ensures an inflammasome response irrespective of the toxins’ substrate specificity.

### TcdB permeabilizes the endolysosomal compartment to ions

Our data demonstrated that LCT-mediated NLRP3 activation required endosomal acidification, indicating that TcdB uptake and trafficking to the late endosome/lysosome is required for NLRP3 activation. To confirm this, we used EGA, a small molecule that inhibits trafficking from the early to late endosome ^21^. Indeed, EGA inhibited IL-1β release (Fig. 2A), demonstrating that trafficking to the late endosome/lysosome is required for NLRP3 activation.

**Figure 2.**
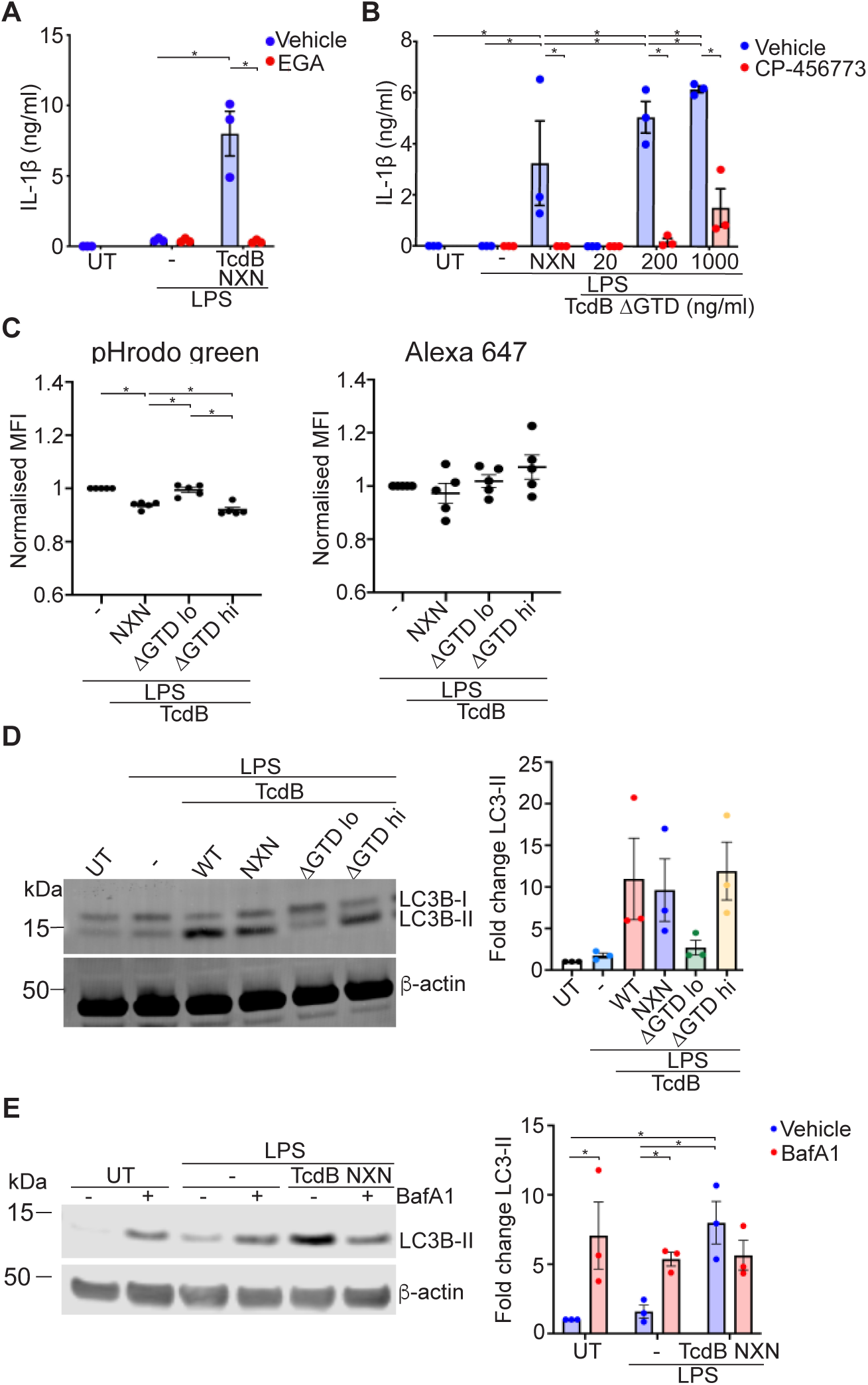
TcdB increases the ion permeability of the endo/lysosomal membrane. **A)** LPS-primed hMDM pre-incubated with EGA (5 μM, 30 min) and treated with TcdB NXN (20 ng/ml, 2 h). IL-1β was measured from sup. **B)** LPS-primed hMDM pre-incubated with CP-456,773 (2.5 μM, 15 min) and incubated with TcdB NXN (20 ng/ml, 2 h), or with increasing concentrations of TcdB ΔGTD (2 h). IL-1β was measured from sup. **C)** MFI of pHrodo green- or A647-coupled 10 kDa dextrans from LPS-primed hMDM treated with TcdB mutants (TcdB NXN and TcdB ΔGTD lo: 20 ng/ml, TcdB ΔGTD hi: 1 μg/ml, 2 h). MFI was normalized to LPS alone. **D)** Immunoblot labeling LC3B or actin in LPS-primed hMDM treated with the different TcdB mutants (see C) for 45 min. LC3B-II was quantified by densitometry. **E)** LPS-primed hMDM treated with TcdB NXN and BafA1 (200 nM, 15 min), which was included 30 min post addition of TcdB NXN. LC3B-II was quantified by densitometry. Immunoblots representative of n = 3. Statistical analysis performed using a two-way ANOVA (A-B), (E) or one-way ANOVA (C-D). *p<0.05, only significant differences annotated.

It has been previously shown that insertion of TcdB into a lipid-bilayer membrane permeabilised the membrane to monovalent cations ^18^. Interestingly, at a 20-fold higher concentration compared to TcdB WT, the TcdB ΔGTD mutant also triggered ion translocation across a lipid-bilayer compared ^18^. Thus, we hypothesized that TcdB ΔGTD activates NLRP3 at higher concentrations that are required to increase the ion permeability of the endolysosomal membranes. To determine this, we incubated LPS-primed hMDM with TcdB ΔGTD at 20 ng/ml (normal concentration for TcdB experiments), 200 ng/ml or 1 μg/ml. TcdB ΔGTD triggered both IL-1β release (Fig. 2B) and caspase-1 cleavage (Fig. S2A) only at concentrations > 20 ng/ml in a concentration- and NLRP3-dependent manner, indicating that an increase in ion permeability of the endolysosomal systems underlies NLRP3 activation.

To determine whether TcdB perturbed endolysosomal ion homeostasis, we assessed lysosomal acidification using fluorescent dextrans conjugated to pHrodo green to monitor pH or to Alexa fluor 647 (A647) as a non-pH sensitive dye control. Both dextrans showed a vesicular pattern with overlapping signals, demonstrating that they localize to the same compartment (Fig. S2B). We then treated LPS-primed hMDM with TcdB NXN, as well as TcdB ΔGTD hi (1 μg/ml) or TcdB ΔGTD lo (20 ng/ml). TcdB NXN was used for this experiment and subsequent experiments to avoid the confounding effects caused by the inhibition of RhoA/B/C, Rac2/3, and Cdc42. We also included the NLRP3 inhibitor CP-456,773 in all conditions to ensure that any effects we observed were not the result of effects downstream of NLRP3 inflammasome activation. Flow cytometry analysis demonstrated that both TcdB NXN and TcdB ΔGTD hi decreased the mean fluorescence intensity (MFI) of pHrodo green, representing a higher pH, whereas TcdB ΔGTD lo had no effect (Fig. 2C). Furthermore, the MFI of A647 was not changed by any of the treatments (Fig. 2C). These results demonstrate that TcdB reduces lysosomal acidification at a concentration that increases ion membrane permeability.

To further confirm this finding, we assessed the non-canonical autophagic response (CASM). CASM is triggered by an increase of the pH in the lumen of endo/lysosomes, most often caused by proton leakage ^22^. This results in conjugation of LC3-I to lipids in the outer leaflet of the endolysosomal membrane, converting LC3-I to its lipidated form, LC3-II. We hypothesized that TcdB would trigger CASM by enabling efflux of protons and/or cations from the lumen of endo/lysosomes. Indeed, TcdB WT, TcdB NXN, and TcdB ΔGTD hi, which all activate NLRP3, also increased LC3B-II in LPS-primed hMDM (Fig. 2D), while TcdB ΔGTD lo had no effect (Fig. 2D), suggesting that TcdB triggered CASM. CASM differs from canonical autophagy as it requires recruitment of the V_1_ subunit of the V-ATPase to the V_0_ subunit on the endolysosomal membrane, which is inhibited by BafA1 ^23^. To determine whether the LC3B-II increase was due to CASM, we incubated LPS-primed hMDM with BafA1 20 min post-addition of TcdB NXN, demonstrating that BafA1 reduced the TcdB NXN-dependent increase in LC3-II (Fig. 2E).

One possible outcome of the TcdB-mediated increase in ion membrane permeability could be a loss of lysosomal integrity, which has been previously suggested to be involved in NLRP3 activation ^1,24^. To determine whether TcdB caused a loss of lysosomal integrity, we generated a BLaER1 human macrophage cell line expressing a galectin-8-mCitrine fusion protein, which binds to carbohydrates in the lumen of damaged endosomes and forms puncta ^25^. We treated these cells with TcdB NXN and investigated puncta formation. However, TcdB NXN did not elicit galectin puncta formation (Fig. S2C), whereas LeuLeu-methyl ester, which causes lysosomal damage, triggered formation of large numbers of puncta (Fig. S2C). This demonstrates that TcdB does not cause loss of lysosomal integrity, which is in line with previous reports, showing an absence of galectin-3 or −8 puncta in TcdB-treated Hep-2 cells ^19^.

### TcdB triggers an increase in cytosolic sodium to activate NLRP3

Our findings indicated that ion translocation from the endolysosomal system to the cytosol is sufficient to activate NLRP3. In fact, NLRP3 activation has been previously associated with a loss of ion homeostasis ^11^. Of the ions that are at a much higher concentration in the endolysosomal system compared to the cytosol, we rationalized that sodium was the most plausible candidate, as it plays an important role in cellular ion homeostasis. To test whether TcdB increased cytosolic sodium, we loaded hMDM with ION Natrium Green-2 (ING-2), a dye that increases in fluorescence when the sodium concentration increases, and measured the change of fluorescence intensity across time in individual cells in response to TcdB NXN. To rule out any confounding effects due to NLRP3 activation, hMDM were incubated with CP-456,773 prior to addition of TcdB NXN. TcdB NXN, but not LPS alone, evoked an increase in ING-2 fluorescence within 15 min after its addition (Fig. 3A-C, Video S1-4). The increase was inhibited by pre-incubation with BafA1 (Fig. 3A-C, Video S1-4), demonstrating that acidification of the endolysosomal compartment was required for the TcdB NXN mediated increase in cytosolic sodium. We confirmed that the increase in cytosolic sodium correlated with the effect of TcdB on ion permeability of the endolysosomal membrane as only TcdB ΔGTD hi, but not TcdB ΔGTD lo increased ING-2 fluorescence (Fig. 3D). The TcdB-driven increase in cytosolic sodium temporally correlated with TcdB-mediated modification of Rac2/3, which also occurred within 15 min, indicating that the toxin reaches the late endosomes/lysosomes within this time period (Fig. S3A). To quantify the amount of sodium translocating to the cytosol following incubation with TcdB, we generated a standard curve by incubating ING-2-loaded hMDM in a balanced salt solution containing different concentrations of sodium and gramicidin, a K^+^/Na^+^ ionophore, to equilibrate the extracellular and intracellular sodium concentrations (Fig. 3E). Based on this curve, the cytosolic sodium concentration prior to addition of TcdB NXN (0 min) was approximately 38 mM at a normalized MFI (F/F0) of 1.433 (Fig. 3E). Addition of TcdB NXN increased the fluorescence signal to maximal, as it could not be increased further by the addition of gramicidin (Fig. 3E). Thus, TcdB resulted in an at least 60 mM increase in the cytosolic sodium concentration. To investigate the contribution of TcdB-mediated sodium increase in the cytosol to the total cellular Na^+^ concentration, we performed atomic absorption spectroscopy (AAS). Compared to LPS alone, TcdB NXN triggered an increase in the overall amount of Na^+^ in the cell (Fig. S3B), demonstrating that TcdB increased cellular sodium levels.

**Figure 3.**
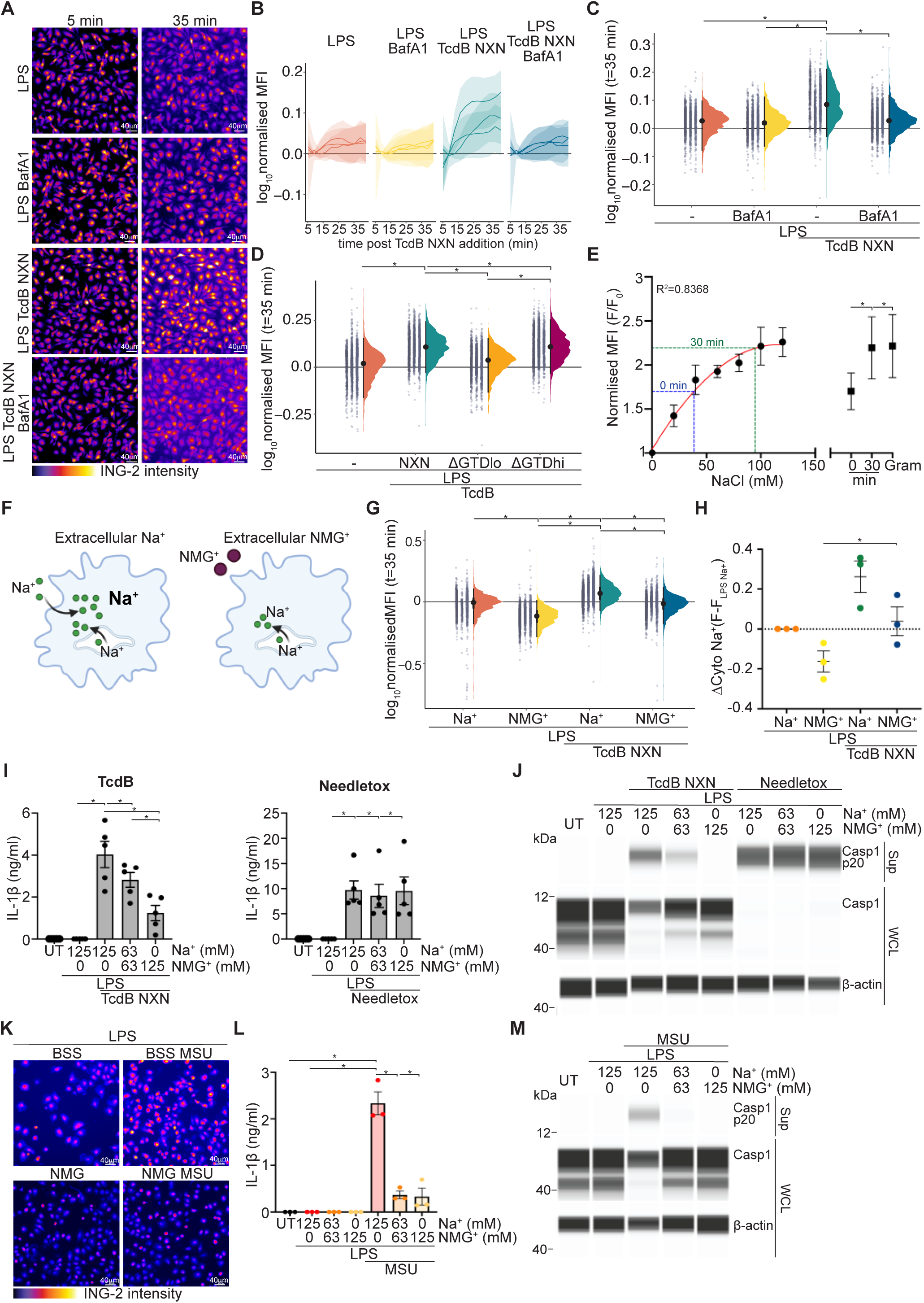
NLRP3 activation by TcdB and MSU crystals is dependent on an acute increase in cytosolic sodium. **A)** Fluorescent images of LPS-primed, ING-2 loaded hMDM stimulated with TcdB NXN and BafA1. **B)** Temporal changes of ING-2 intensity normalised to t = 0 min, per cell. Mean (solid line) ± S.D. (shaded area) is shown for each experimental replicate. MFI = Mean Fluorescence Intensity. **C)** Normalized ING-2 intensities at t = 35 min from B). Grey dot represents one cell, mean ± S.D. are shown. **D)** Normalised ING-2 intensity per cell for LPS-primed hMDM with TcdB NXN or TcdB ΔGTD lo (20 ng/ml) or hi (1 μg/ml). **E)** Normalized ING-2 MFI from LPS-primed hMDM incubated with different extracellular [Na^+^] in the presence of gramicidin for 10 min (circles) or LPS-primed hMDM incubated with TcdB NXN imaged at 0 and 30 min and after addition of gramicidin (squares, >400 cells/concentration). Mean ± SEM for n = 3 is shown. **F)** Possible sources of cytosolic Na^+^, NMG^+^ substitution removes extracellular Na^+^. **G)** Normalized ING-2 MFI from LPS-primed hMDM incubated with TcdB NXN in Na^+^ or NMG^+^ BSS. **H)** Mean ΔING-2 MFI for each condition from G). Mean ± SEM is shown, each point represents one donor/experimental repeat. LPS-primed hMDM incubated with TcdB NXN or Needletox (25 ng/ml, 2 h) in Na^+^ or NMG^+^ containing BSS. IL-1β was assessed from sup **(I)** or caspase-1 was assessed from sup or WCL by SWA **(J)**. **K)** Fluorescence images of LPS-primed, ING-2 loaded hMDM stimulated with MSU crystals (100 μg/ml) in Na^+^ or NMG^+^ containing BSS. LPS-primed hMDM incubated with MSU in Na^+^ or NMG^+^ containing BSS. IL-1β was assessed from sup **(L)** or caspase-1 was assessed from sup or WCL by SWA **(M)**. Mean ± SEM n = 3 – 5 shown for I and L, each point represents one donor. SWAs representative of n = 2. (A-H) and (K) were performed in the presence of VX-765. Statistical analysis performed using one-way ANOVA *p<0.05, only significant differences annotated.

We also assessed whether TcdB had any effect on the Na^+^/K^+^ ATPase, which would be expected to remove sodium from the cytosol. However, using rubidium uptake assays, where Rb^+^ serves as a non-naturally occurring tracer for Na^+^/K^+^ ATPase-dependent K^+^ transport ^26^, we found no significant differences in Rb^+^ levels after TcdB addition, whereas treatment of cells with ouabain resulted in a decrease in Rb^+^. These findings demonstrate that inhibition of the Na^+^/K^+^ ATPase did not contribute to TcdB-driven Na^+^ accumulation in the cytoplasm (Fig. S3C).

To determine whether the TcdB-mediated increase in cytosolic sodium was required for NLRP3 activation, we stimulated LPS-primed hMDM with TcdB in a balanced salt solution (BSS), substituting the extracellular Na^+^ with N-methyl-D-glucamine chloride (NMG^+^), a large monovalent cation that maintains osmolarity and membrane potential, but is inefficiently transported by cellular pumps and ion channels ^27^. Importantly, replacing Na^+^ BSS with NMG^+^ BSS immediately removes extracellular Na^+^ but not endolysosomal Na^+^, which will only be exchanged slowly as NMG^+^ is endocytosed by the cell (Fig. 3F). We first assessed whether substituting extracellular Na^+^ with NMG^+^ reduced the increase in cytosolic Na^+^. Consistent with Na^+^ entering the cytosol from both endolysosomal and extracellular compartments, substitution of extracellular Na^+^ with NMG^+^ reduced, but did not ablate the TcdB-driven in-crease in cytosolic sodium (Fig. 3G, S3D, Video S5-8). In the presence of ex-tracellular NMG^+^ the increase in the cytosolic sodium concentration was sig-nificantly higher upon treatment with TcdB NXN than with LPS alone (Fig. 3H) but did not reach the same concentration in the presence of extracellular Na^+^ (Fig. 3H). This indicates that sodium translocation from the endolysosomal system to the cytosol is the initiating event triggering cytosolic sodium accumulation.

Preventing the increase in cytosolic Na^+^ by NMG^+^ substitution inhibited TcdB-dependent NLRP3 activation in a concentration-dependent manner, as meas-ured by IL-1β release (Fig. 3I) and caspase-1 cleavage (Fig. 3J) but did not affect Needletox-dependent IL-1β release (Fig. 3I) or caspase-1 cleavage (Fig. 3J) mediated by NLRC4. NMG^+^ substitution did not prevent TcdB-mediated modification of Rac2/3 (Fig. S3E), demonstrating that substitution of Na^+^ with NMG^+^ inhibited NLRP3 activation downstream of TcdB insertion into the endolysosomal membrane. This collectively demonstrates that the TcdB-mediated increase in cytosolic sodium activates the NLRP3 inflammasome.

### NLRP3 activation by particulate matter requires an increase in cytosolic sodium

Our data demonstrated that translocation of sodium from the endocytic system to the cytosol activates NLRP3. Other noxious molecules, including MSU and silica crystals, have been suggested to activate NLRP3 in a cathepsin-dependent manner by perturbing endo/lysosomes ^1^. However, a recent study using a combined knock-out of 5 cathepsins only partially decreased NLRP3 activation caused by lysosomal rupture ^28^. Therefore, we hypothesized that particulate matter also increases the ion permeability of the endolysosomal compartment, thereby activating NLRP3. We first confirmed that MSU-mediated NLRP3 activation required phagocytosis in hMDM (Fig. S3F). Next, we assessed cytosolic Na^+^ levels in hMDM incubated with MSU in the presence or absence of extracellular Na^+^. Similar to TcdB, incubating hMDM with MSU triggered an increase in ING-2 fluorescence (Fig. 3K, S3G-H), that was absent in the presence of NMG^+^ (Fig. 3K, S3G-H), demonstrating _that_ MSU also increases the cytosolic sodium concentration. We then investigated whether the MSU-dependent increase in cytosolic sodium was required for NLRP3 activation. Replacing Na^+^ by NMG^+^ prevented MSU-dependent IL-1β release (Fig. 3L) and caspase-1 cleavage (Fig. 3M), demonstrating that it was sufficient to prevent activation of NLRP3. Furthermore, treatment of hMDM with silica crystals also triggered an increase of the cytosolic sodium concentration (Fig. S3I-K), which was required for NLRP3 activation (Fig. S3L), though to a lesser degree than MSU. These results demonstrate that translocation of sodium from the endolysosomal system to the cytosol is a common requirement for activation of NLRP3 by particulate matter.

### TcdB-mediated NLRP3 activation requires macropinocytosis

Our results established that, in addition to translocation of endolysosomal sodium, entry of extracellular sodium into the cytosol was also required for TcdB-mediate NLRP3 activation. One mechanism of sodium entry into the cell is macropinocytosis, a specialized form of endocytosis that is characterized by the non-specific uptake of large amounts of extracellular fluid, which contains high sodium concentrations ^29^. hMDM perform macropinocytosis prolifically and, unlike other cells, constitutively ^30^. Therefore, we targeted different processes required for macropinosome formation using pharmacological inhibitors: EIPA, which inhibits Na^+^/H^+^ exchange, LY-294002, a class I PI3K inhibitor, two inhibitors of actin rearrangement, jasplakinolide and latrunculin B, and NPS2143, an inhibitor of the calcium sensing GPCR (CaSR), which inhibits constitutive macropinocytosis in hMDM ^30^ (Fig. 4A). As anticipated, all compounds inhibited macropinocytosis, as measured by uptake of a 70 kDa Oregon green-conjugated dextran (Fig. S4A). Importantly, all compounds inhibited TcdB-mediated NLRP3 activation as measured by IL-1β release (Fig. 4B), but did not inhibit activation of NLRP3 by nigericin (Fig. 4B). Moreover, they did not inhibit entry of TcdB into the cytoplasm, as measured by modification of Rac2/3 (Fig. S4B). We confirmed these results by knocking down the CaSR (Fig. S4C), which reduced macropinocytosis (Fig. S4D), and inhibited NLRP3 activation by TcdB (Fig. 4C), but had no effect on Needletox-dependent activation of NLRC4 (Fig. 4C). These results demonstrate that macropinocytosis is required for TcdB-mediated NLRP3 activation.

**Figure 4.**
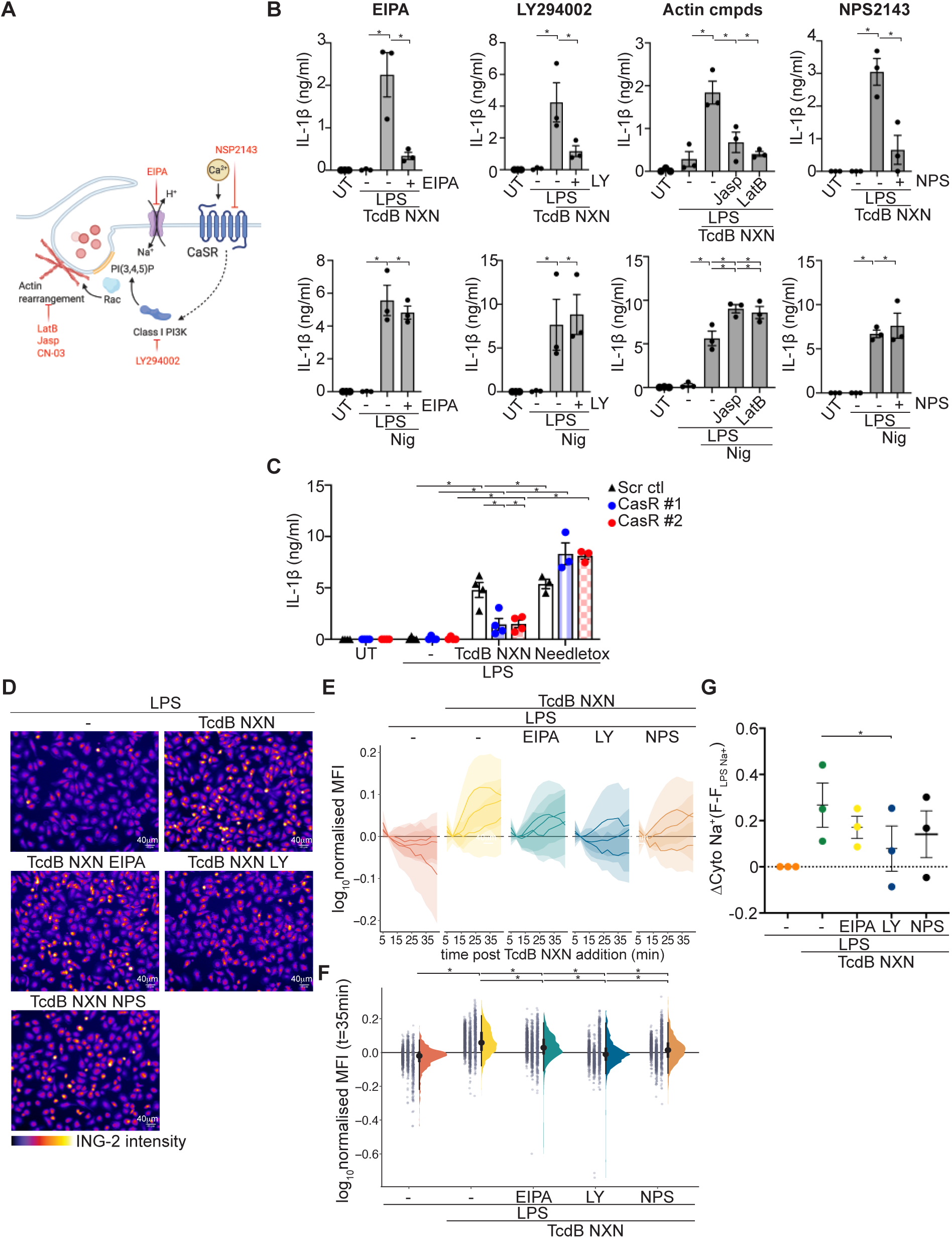
TcdB-mediated NLRP3 activation and cytosolic sodium accumulation requires macropinocytosis. **A)** The macropinocytosis inhibitors used and their targets. LPS-primed hMDM pre-incubated with EIPA (20 μM), LY294002 (10 μM), Jasplakinolide (0.5 μM), Latrunculin B (2 μM), or NPS2143 (5 μM) for 30 min, then treated with either TcdB NXN (20 ng/ml) (upper row) or nigericin (8 μM) (bottom row) for 2h. IL-1β was assessed from sup **(B). (C)** LPS-primed hMDM electroporated with CaSR targeting or scrambled (scr) siRNA treated with TcdB NXN (20 ng/ml) or needletox (25 ng/ml). IL-1β assessed from the sup. **D)** Fluorescent images of LPS-primed, ING-2 loaded hMDM stimulated with TcdB NXN +/- macropinocytosis inhibitors as listed above at t = 30 min. **E)** Temporal changes of ING-2 intensity normalised to t = 0 min, per cell. Mean (solid line) ± S.D. (shaded area) is shown for each experimental replicate. **F)** Normalized ING-2 intensities at t = 35 min from B). Grey dot represents one cell, mean ± S.D. are shown. **G)** Mean ΔING-2 MFI for each condition from G). Mean ± SEM, n = 3 – 4, each point represents one donor. (D-G) were performed in the presence of VX-765. Statistical analysis performed using one-way ANOVA *p<0.05, only significant differences annotated.

To test whether inhibition of macropinocytosis prevented the TcdB-driven increase in cytosolic sodium, we pre-incubated hMDM with the macropinocytosis inhibitors, and then assessed changes in the cytosolic Na^+^ concentration following addition of TcdB NXN. Inhibiting macropinocytosis reduced both the velocity and amplitude of the TcdB-mediated increase in the cytosolic Na^+^ concentration (Fig. 4D-E, Video S9-13), resulting in a decrease in the mean fluorescence at 35 mins post addition of TcdB NXN (Fig. 4F-G). However, inhibiting macropinocytosis did not reduce ING-2 intensity to the level of hMDM treated with LPS alone (Fig. 4G), demonstrating that, consistent with our hypothesis, macropinocytosis reduces the TcdB-mediated increase in cytosolic Na^+^ but does not ablate it. To determine if macropinocytosis contributed the TcdB uptake, we inhibited macropinocytosis and assessed TcdB uptake by immunofluorescence. However, with the exception of latrunculin B, inhibition of macropinocytosis did not affect TcdB uptake (Fig. S4E) and also did not change CASM as measured by LC3B-II levels, indicating that that these inhibitors do not change the amount of TcdB inserting into the endolysosomal membrane (Fig. S4F). These results demonstrate that macropinocytosis is required for TcdB-driven NLRP3 activation by contributing to TcdB-mediated cytosolic sodium accumulation.

### The TcdB-mediated increase in cytosolic sodium activates NLRP3 by triggering changes in cell volume

A key role of cellular sodium is to function as an osmolyte to regulate cell volume. Previous studies have also linked dysregulation of cell volume to NLRP3 activation ^31,32^, suggesting that the increase in cytosolic sodium may activate NLRP3 through the same mechanism. Therefore, we hypothesized that the TcdB-mediated increase in cytosolic sodium activated NLRP3 by promoting water influx to trigger cell swelling. To this end, we measured the cell volume of hMDM following incubation with TcdB NXN using a Coulter counter. TcdB NXN increased the mean cell volume of the hMDM population (Fig. 5A) as well as increasing the percentage of hMDM with a cell volume greater than the median of the control hMDM (Fig. 5B). This was evident at 30 min and peaked at 45 min (Fig. 5A-B). Similarly, TcdB ΔGTD hi, but not TcdB ΔGTD lo increased cell volume 30 min post addition (Fig. 5C), correlating with cytosolic sodium accumulation and NLRP3 activation. We next determined whether the TcdB-dependent increase in cell volume correlated with the increase caused by incubation in hypo-osmotic media, which was previously shown to activate NLRP3 ^31^. We first confirmed that hypo-osmotic medium triggered IL-1β release from LPS-primed hMDM (Fig. S5A). Addition of hypo-osmotic medium increased the peak mean cell volume (Fig. S5B), though to a lesser degree than TcdB, and it increased the proportion of cells with a cell volume greater than the median (Fig. S5C), demonstrating that the TcdB-mediated volume increase is comparable to a volume increase known to activate NLRP3.

**Figure 5.**
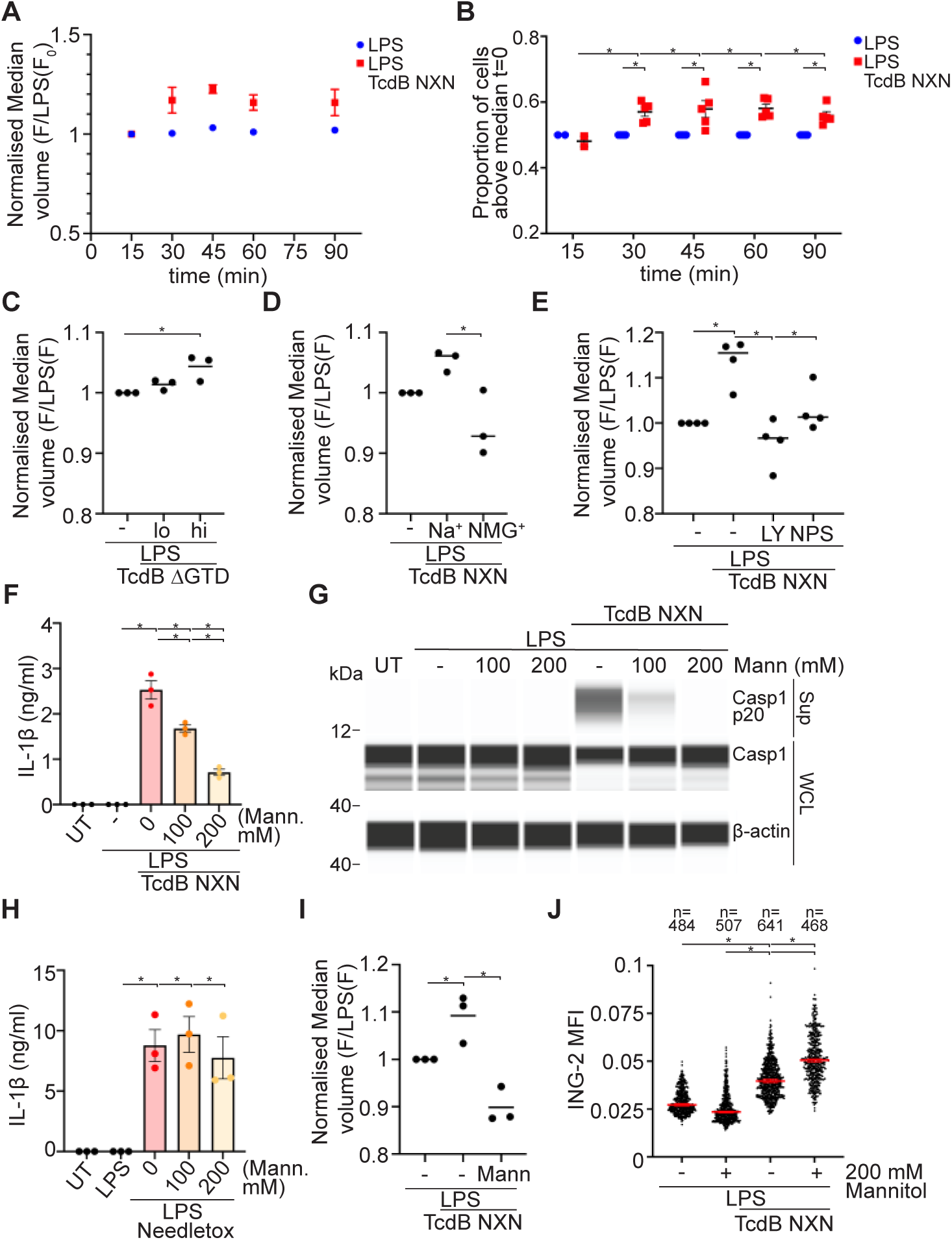
The TcdB-mediated increase in cytosolic sodium activates NLRP3 by triggering cell swelling. **A)** Normalised median cell volume of LPS-primed hMDM incubated with TcdB NXN or **B)** the proportion of hMDM greater than the median. Mean ± SEM, n = 5. **C)** Normalized median cell diameter (30 min) of LPS-primed hMDM incubated with TcdB ΔGTD lo (20 ng/ml) or hi (1 μg/ml). Mean, n = 3. **D)** Normalised median cell diameter (30 min) of LPS-primed hMDM +/- TcdB NXN in Na^+^ or NMG^+^ BSS. Mean, n = 3. **E)** Normalised median cell diameter for LPS-primed hMDM with either LY294002 or NPS2143 and VX-765, analysed after 40 min TcdB NXN. Mean, n = 4. LPS-primed hMDM incubated with TcdB NXN in media containing 100 mM or 200 mM mannitol. IL-1β was assessed from sup **(F)** or caspase-1 was assessed from sup or WCL by SWA, representative of n = 2 **(G)**. LPS-primed hMDM incubated with needletox in media containing 100 mM or 200 mM mannitol. IL-1β was assessed from sup **(H)**. **I)** Normalized median cell diameter (30 min) of LPS-primed hMDM +/- TcdB NXN in media +/- 200 mM mannitol. Mean, n = 3. **J)** MFI of ING-2 from LPS-primed hMDM +/- TcdB NXN +/- 200 mM mannitol at t = 30 min. Median ± 95% CI are shown. For F) and H), Mean ± SEM, n = 3. For all experiments each point represents one donor. (A-E and I) were performed in the presence of VX-765. Statistical analysis performed using one-way ANOVA *p<0.05, only significant differences annotated.

To investigate whether cytosolic sodium accumulation was required for TcdB-driven cell swelling, we incubated hMDM with TcdB in the presence or absence of Na^+^. Substitution of Na^+^ with NMG^+^ prevented the TcdB-driven cell swelling (Fig. 5D). Similarly, blocking the TcdB-mediated cytosolic sodium increase by pre-incubating hMDM in macropinocytosis inhibitors, both of which inhibited the TcdB-driven increase in cytosolic sodium, prevented the cell volume increase triggered by TcdB (Fig. 5E). This demonstrates that the TcdB-mediated increase in cell volume was driven by the increase in cytosolic sodium.

To determine whether the osmolarity-driven, TcdB-mediated cell-volume increase activates NLRP3, we incubated LPS-primed hMDM with TcdB in extracellular media containing an additional 100 mM or 200 mM mannitol. This increases the osmolarity of the extracellular medium, thereby inhibiting cell swelling by preventing water influx into the cytosol. Addition of mannitol inhibited TcdB-mediated activation of NLRP3 in a concentration dependent manner, as measured by IL-1β release (Fig. 5F) and caspase-1 cleavage (Fig. 5G). In contrast, mannitol did not affect NLRC4 inflammasome-dependent IL-1β release following activation by needletox (Fig. 5H). Importantly, mannitol prevented the TcdB-mediated increase in cell volume (Fig. 5I), but did not alter the TcdB-driven increase in cytosolic sodium (Fig. 5J), cellular sodium, (Fig. S5D) or TcdB uptake and internalization (Fig. S5E). These results demonstrate that the TcdB-driven increase in cytosolic sodium activated NLRP3 through cell volume change.

### Increased cytosolic sodium perturbs osmotic regulation of endocytic trafficking to enable TcdB-mediated NLRP3 activation

In addition to its effects on cellular volume, sodium also regulates the osmotic balance of the endolysosomal system, in particular endocytic trafficking ^33^. Notably, endocytic trafficking is regulated by the endocytic membrane tension, which is controlled by the passive efflux of sodium, chloride, and subsequently water from the endosomal lumen into the cytosol (Fig. S6A). Recent studies suggested that inhibition of endosomal trafficking is required for NLRP3 activation ^12,34^, although the underlying molecular mechanisms remain unclear. We hypothesized that the TcdB-mediated increase in cytosolic sodium subsequently inhibits endocytic trafficking by preventing sodium efflux from endosomes, thus preventing the subsequent drop in endosomal membrane tension, inhibiting endocytic trafficking and, thereby, activate NLRP3.

We first assessed whether TcdB inhibited endosomal trafficking using a pulse-chase assay with A647-labelled transferrin (Tfn-A647). Trafficking of the lig-and-bound Tfn receptor complex is clathrin-dependent; the receptor is transported to the early endosome following internalization before being recycled back to the plasma membrane (Fig. S6B). As such, inhibition of endocytic trafficking increases the number of Tfn-A647-positive vesicles. To determine if endocytic trafficking was inhibited by TcdB NXN, we pulsed LPS-treated hMDM for 10 min with Tfn-A647 at 25 min post TcdB NXN treatment, when TcdB NXN already triggered an increase in cytosolic sodium (see Fig. 3B). We then determined the number of cells with Tfn-A647-positive (Tfn^+^) vesicles at 10 min intervals (Fig. 6A). Both LPS- and LPS and TcdB NXN-treated hMDM contained Tfn+ vesicles following the 10 min pulse (0 min) as anticipated (Fig. 6A). However, only the TcdB NXN-treated hMDM still had a proportion of hMDM with Tfn^+^ vesicles even after a 30 min chase period (Fig. 6A), indicating that TcdB inhibited endocytic trafficking. Tfn trafficking was similarly inhibited in hMDM treated with TcdB ΔGTD hi but not TcdB ΔGTD lo, correlating with TcdB-mediated cytosolic sodium increase and NLRP3 activation (Fig. S6C). Tfn+ vesicles in the TcdB-treated hMDM colocalized with Rab5, a marker of the early endosome (Fig. 6B). To confirm that the increase in Tfn^+^ hMDM was due to inhibition of endocytic trafficking and not differential Tfn uptake, we assessed Tfn-A647 binding. LPS-primed hMDM treated with TcdB NXN for 25 min were incubated with Tfn-A647 on ice for 1 h, then incubated at 37°C for 1 min to enable Tfn-647 internalisation, then washed to remove any residual Tfn-A647. Quantification by immunofluorescence demonstrated that TcdB did not alter Tfn-A647 binding (Fig. S6D).

**Figure 6.**
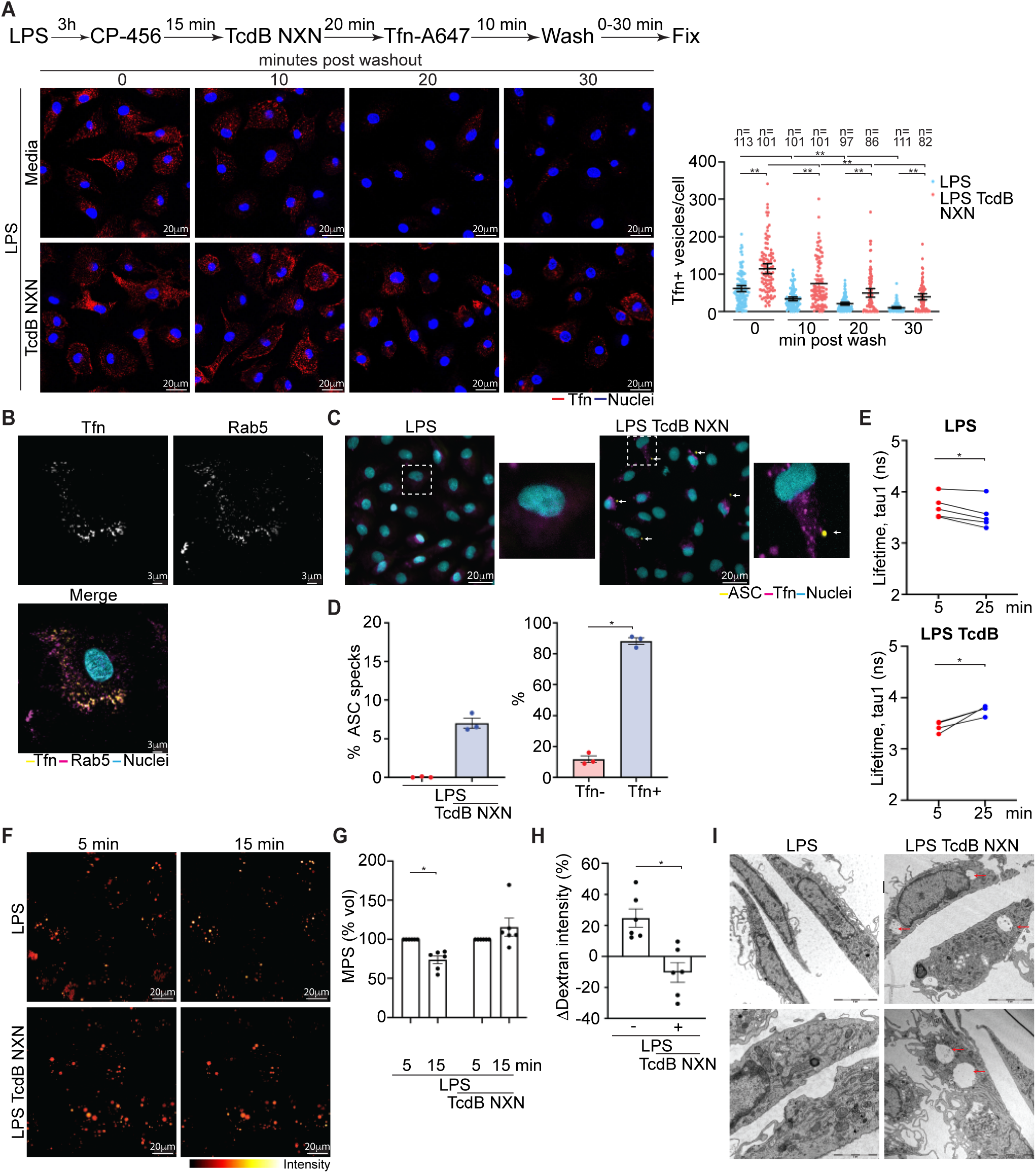
TcdB inhibits osmotic resolution of endosomes to enable NLRP3 inflammasome activation. **A)** Fluorescence confocal images and quantitation of Tfn-A647 pulse-chase in LPS-primed hMDM treated with TcdB NXN. Mean ± 95% CI are shown, each dot represents one cell, data is from pooled from 3 experimental replicates. **B)** Fluorescence confocal images of Tfn-A647 and Rab5 labeling in LPS-primed-hMDM treated with TcdB NXN. Representative of 50 cells per donor, n = 3. **C)** Fluorescence images and **D)** quantitation of ASC-speck^+^ and Tfn^+^ hMDM LPS-primed, pre-incubated with VX-765 and treated with TcdB NXN. Mean ± SEM, n = 3, approx. 1500 cells analyzed/donor. **E)** FLIM analysis of LPS-primed, EE-flipper (2 μM) loaded hMDM treated with TcdB NXN, n = 3-5 images, 2-3 cells per image, mean shown. **F)** Fluorescence confocal images of LPS-primed hMDM treated with TcdB NXN pulsed with 70 kDa Oregon green (OG) labelled dextran and 200 U/ml M-CSF for 4 min, then washed and imaged over time. **G)** Macropinosome volume measurement at t = 5 and t = 15 min. **H)** ΔDextran intensity for macropinosomes between t = 5 and t = 15 min. Each dot represents the average volume and intensity for an experimental repeat, mean ± SEM, n = 6. **I)** Transmission electron microscopy image of LPS-primed hMDM incubated with TcdB NXN (45 min), representative of 3 independent donors. For all experiments except D), hMDM were pre-incubated with CP-456,773. Statistical analysis performed with two-way ANOVA (A) or Student’s t-test (D-E and G-H). *p<0.05, only significant differences annotated.

To confirm that TcdB generally inhibited endocytic trafficking, we assessed endocytic transport of Alexafluor-555-labeled cholera toxin subunit B (CtxB). In contrast to Tfn, CtxB trafficking is clathrin-independent, and is transported to the cytosol rather than being re-externalized (Fig. S6E). Incubation with TcdB NXN increased the proportion of hMDM with CtxB-positive vesicles (CtxB^+^) compared to LPS-primed hMDM (Fig. S6F). This was not due to differential uptake of CtxB, as the overall amount of CtxB taken up, as measured by the integrated intensity of the cell, was unchanged by TcdB (Fig. S6F). CtxB^+^ vesicles colocalized with Rab5 (Fig. S6G). These results demonstrate that TcdB inhibits endosomal trafficking at the level of the early endosome, which correlates with both an increase in cytosolic sodium and NLRP3 activation.

Our results revealed a strong association between TcdB-mediated inhibition of endocytic trafficking and NLRP3 activation. To confirm this, we correlated trafficking inhibition with a marker of inflammasome activation, ASC-speck formation ^35^. Unlike other inflammasome readouts, ASC speck formation can be measured on a single cell level, and so can be correlated with cells with Tfn^+^ vesicles. Only LPS-primed hMDM incubated with TcdB NXN displayed ASC specks (Fig. 6C), with around 10% of the population responding (Fig. 6D). Of these, almost all hMDM with an ASC speck were Tfn^+^ (Fig. 6D), demonstrating a strong correlation between NLRP3 activation and endosomal trafficking inhibition.

As we established that TcdB inhibited endocytic trafficking, we next hypothesized whether the increase in cytosolic sodium prevented a loss of endosomal membrane tension. To test this, we assessed endocytic membrane tension using the early endosomal flipper dye (EE-flipper), which integrates into endosomal membranes and changes fluorescence lifetime based on endosomal membrane tension ^36^. We first measured EE flipper fluorescence lifetime changes in response to mannitol, which decreased (Fig. S7A), and monensin, which increased endosomal membrane tension (Fig. S7B). We then assessed whether TcdB altered endosomal membrane tension in LPS-primed hMDM and found that incubation with TcdB NXN consistently resulted in an increase in fluorescence lifetime, indicative of increased membrane tension (Fig. 6E, S7C). In contrast, fluorescence lifetime decreased in the hMDM incubated with LPS alone (Fig. 6E, S7C).

Having established that TcdB indeed leads to an increase in endosomal membrane tension, we next tested whether this was due to sodium efflux from the endosomes. To this end, we assessed macropinosome volume as a surrogate marker. As macropinosomes mature, they decrease in volume, which requires water loss. This water loss is driven by the passive efflux of sodium from the lumen of the macropinosome into the cytosol ^37^. Thus, macropinosomes will be unable to reduce in size when changing the sodium gradient through increasing cytosolic sodium, which prevents passive sodium efflux from macropinosomes into the cytosol. To first stimulate macropinosome formation in hMDM, we pulsed them with M-CSF in the presence of a 70 kDa fluorescent dextran for 5 min. The dextran fluorescence intensity can be used as a read-out for macropinosome volume as the dextran gets concentrated in the macropinosome lumen when they reduce in volume, whereby the fluorescence intensity increases ^37^. Macropinosomes formed in the LPS-primed hMDM decreased in size (Fig. 6F, S7D, Video S14), while the macropinosomes in the TcdB-treated hMDM remained unchanged (Fig. 6F, S7D, Video S15). Analysis confirmed that the macropinosomes formed in the LPS-primed hMDM decreased in size (Fig. 6G) and increased in fluorescence intensity (Fig. 6H), whereas both the size and fluorescence intensity of the macropinosomes formed in the TcdB-treated hMDM remained constant (Fig. 6G-H), demonstrating that TcdB prevented macropinosome volume reduction. To assess macropinosomes independently of dextran loading, we performed transmission electron microscopy (TEM). LPS-primed hMDM treated with TcdB NXN displayed large vacuolar structures lacking electron density, which were absent in the LPS-primed hMDM only (Fig. 6I), indicating that macropinosomes are not resolved in the presence of TcdB.

To determine if TcdB inhibited the macropinosome volume decrease by preventing water efflux from the lumen of the macropinosomes, we pulsed the dextran-loaded, LPS-primed and TcdB NXN-treated hMDM with hyperosmotic BSS containing an additional 200 mM mannitol, which triggers water movement from both intracellular compartments and the cytosol to the extracellular space. Accordingly, pulsing TcdB-treated hMDM with hyperosmotic BSS caused the macropinosomes to reduce in size (Fig. S7E, Video S16-17), confirming that the macropinosomes failed to shrink due to retention of water in the lumen of macropinosomes. These results collectively demonstrate that TcdB inhibits endosomal trafficking by preventing osmotic resolution of endosomes and macropinosomes.

### Cytosolic sodium accumulation is a general requirement for NLRP3 activation

Many NLRP3 activators, including ionophores, ATP and pore-forming toxins, trigger potassium efflux from the cytosol, and this is considered to be a crucial step in NLRP3 activation ^38^. In contrast to these activators, we found that TcdB-mediated NLRP3 activation was dependent on an increase in cytosolic sodium, and it was unclear if a decrease in cytosolic potassium was also required. To determine whether TcdB triggered cytosolic potassium efflux, we assessed cellular K^+^ in hMDM following incubation with TcdB NXN, in the presence or absence of extracellular Na^+^ to assess the role of cytosolic sodium accumulation. Compared to the LPS control, TcdB NXN triggered a decrease in cellular K^+^ that was partially rescued in the absence of extracellular Na^+^ (Fig. 7A), indicating that TcdB-mediated cellular K^+^ efflux was partially dependent on cytosolic sodium accumulation. However, the TcdB-mediated potassium efflux did not rely on cell swelling induced by hyperosmotic medium (Fig. S8A), demonstrating that the TcdB-mediated K^+^ efflux was partially dependent on the cytosolic sodium increase but did not require cell volume change.

**Figure 7.**
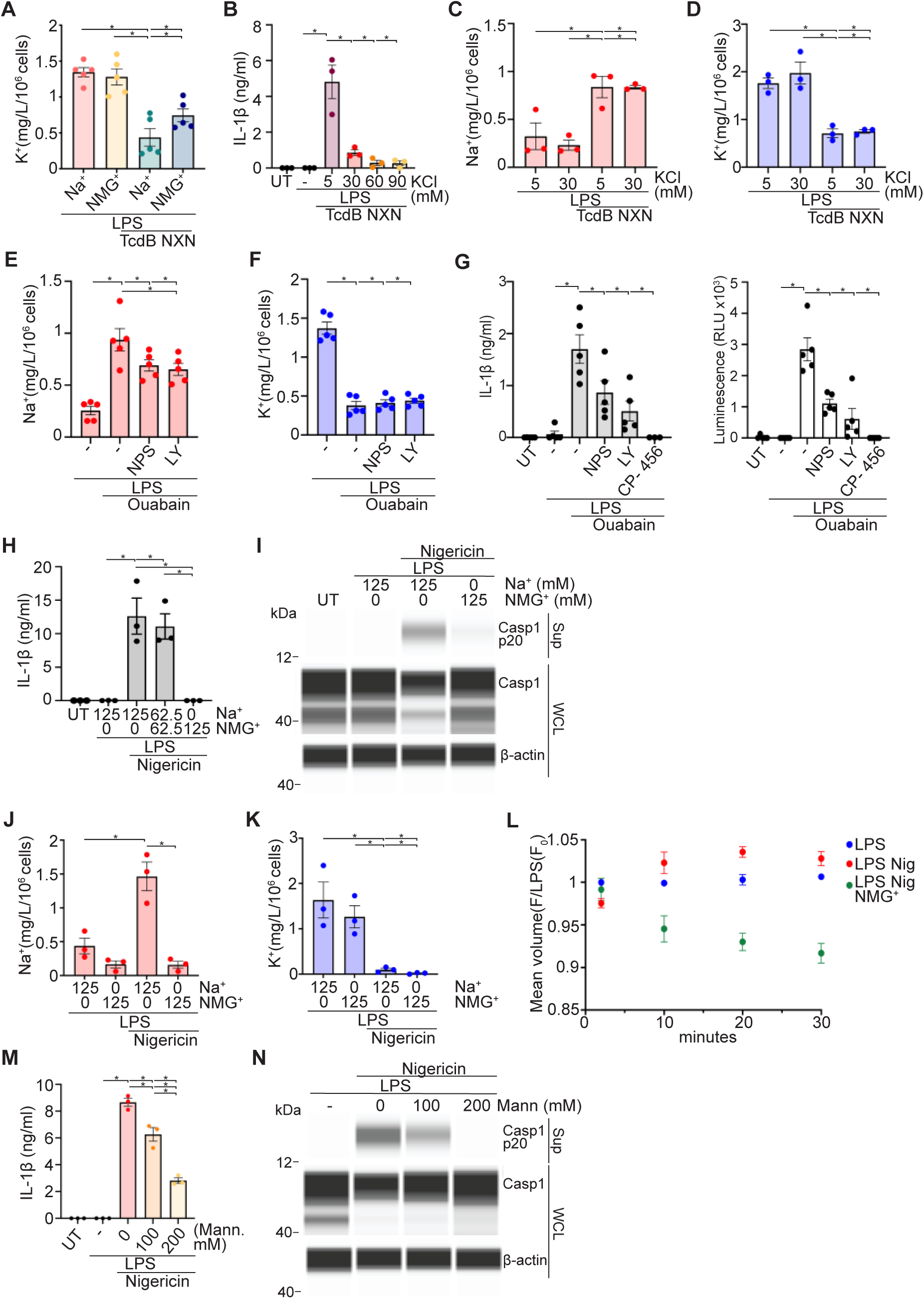
Cytosolic sodium influx is a common requirement for NLRP3 activation. **A)** Cellular K^+^ from LPS-primed hMDM +/- TcdB NXN (20 ng/ml, 45 min) in Na^+^ or NMG^+^ containing BSS. **B)** LPS-primed hMDM incubated with TcdB NXN (20 ng/ml, 2 h) in media containing increasing concentrations of KCl. IL-1β was assessed from sup. Cellular Na^+^ (**C)** and (**D)** K^+^ from LPS-primed hMDM incubated with TcdB NXN (20 ng/ml, 45 min) in media containing 5 mM or 30 mM KCl. Cellular Na^+^ (**E)** and (**F)** K^+^ from LPS-primed hMDM pre-incubated with LY294002 (LY) (10 μM) or NPS2143 (NPS) (5 μM) for 30 min, then treated with ouabain (0.5 mM, 2 h). **(G)** IL-1β and caspase-1 activity were assessed from sup from hMDM treated as in E). LPS-primed hMDM incubated with nigericin (8 μM, 2 h) in Na^+^ or NMG^+^ containing BSS. IL-1β was assessed from sup **(H)**, caspase-1 was assessed from sup or WCL by SWA **(I)**. Cellular Na^+^ **(J)** and K^+^ **(K)** from LPS-primed hMDM incubated with nigericin as in H). **L)** Normalised median cell volume of LPS-primed hMDM incubated with nigericin (8 μM) in Na^+^ or NMG^+^ containing BSS. Mean ± SEM, n = 3. LPS-primed hMDM incubated with nigericin (8 μM, 2 h) in media with 100 mM or 200 mM mannitol. IL-1β was assessed from sup **(M)**, caspase-1 was assessed from sup or WCL by SWA **(N)**. Mean ± SEM, n = 3 – 5, each point represents one donor. SWAs representative of n = 2. A), C)-F) and J)-L) were performed in the presence of VX-765. Statistical analysis performed with one-way ANOVA, *p<0.05, only significant differences annotated.

In previous studies, the requirement for potassium efflux in NLRP3 activation has been assessed by increasing the extracellular K^+^ concentration. In line with this, increasing the extracellular concentration of KCl from 5 mM to 30 mM almost completely ablated TcdB-mediated NLRP3 activation (Fig. 7B). This was due to the increase in extracellular potassium and not chloride as additional extracellular potassium-gluconate inhibited TcdB-mediated NLRP3 activation at the same concentrations as KCl (Fig. S8B). Also, 30 mM KCl did not prevent modification of Rac2/3 by TcdB (Fig. S8C), demonstrating that TcdB uptake and translocation were unaffected. In contrast, NLRC4 inflammasome activation by needletox was unaffected by 30 mM extracellular KCl (Fig. S8D). To determine whether increasing the extracellular K^+^ concentration inhibits the TcdB-driven change in the cellular ion concentrations, we assessed cellular Na^+^ and K^+^ in LPS-primed hMDM incubated with TcdB NXN in media with 5 or 30 mM KCl. However, 30 mM extracellular KCl did not prevent the TcdB-driven increase in cellular Na^+^ (Fig. 7C) and, moreover, did not prevent the decrease in cellular K^+^ (Fig. 7D). Therefore, increasing extracellular K^+^ inhibits TcdB-mediated NLRP3 activation, but this was independent of any effects on cellular Na^+^ or K^+^.

Our data indicated that cytosolic sodium accumulation was the main requirement for NLRP3 activation by TcdB, but it was unclear whether the subsequent decrease in cellular K^+^ was required. This was difficult to resolve using TcdB as the increase in cytosolic Na^+^ was partially required for the decrease in cellular K^+^. Therefore, we used the Na^+^/K^+^ ATPase inhibitor ouabain, which was shown to activate NLRP3 previously ^39^. Unlike TcdB, ouabain should increase cellular Na^+^ and decrease cellular K^+^ independently, as uptake of Na^+^ into the cytosol occurs through different mechanisms to K^+^ efflux. Thus, Na^+^ accumulation in the cytosol can be inhibited without preventing K^+^ efflux. To limit the ouabain-mediated increase in cellular Na^+^, we inhibited macropinocytosis using LY-294002 (LY) or NPS2143 (NPS), which we demonstrated in our previous experiments to decrease the TcdB-driven increase in cytosolic Na^+^ (see Fig. 4E-G). Accordingly, incubating LPS-primed hMDM with ouabain resulted in an increase in cellular Na^+^ (Fig. 7E) and a decrease in cellular K^+^ (Fig 7F). Importantly, inhibiting macropinocytosis only reduced the ouabain-driven increase in cellular Na^+^ (Fig. 7E) but did not alter the decrease in cellular K+ (Fig. 7F). To determine whether this impacted ouabain-mediated NLRP3 activation, we then assessed IL-1β release and caspase-1 activity under the same conditions and found that inhibition of macropinocytosis markedly decreased, but did not ablate, ouabain-mediated NLRP3 activation (Fig. 7G). All IL-1β release and caspase-1 activation triggered by ouabain was NLRP3-dependent, as it was completely inhibited by the NLRP3 inhibitor CP-456,773 (Fig. 7G). These results demonstrate that inhibiting the increase in cytosolic Na^+^ prevents NLRP3 activation even in the presence of cytosolic K^+^ efflux.

In contrast to TcdB and ouabain, many NLRP3 activators, including ionophores such as nigericin, trigger NLRP3 activation by decreasing cellular K^+^ ^38^. However, how the decrease in cellular K^+^ subsequently triggers NLRP3 activation remains an open question. Given the importance of cytosolic sodium accumulation for TcdB-mediated NLRP3 activation, we hypothesized that NLRP3 activators that decrease cellular K^+^ will require a subsequent increase in cytosolic sodium to activate NLRP3. To test this, we incubated LPS-primed hMDM with nigericin, an ionophore that decreases cytosolic K^+^ by exchanging intracellular K^+^ for extracellular H^+^, in the absence or presence of extracellular Na^+^. This revealed that substitution of Na^+^ for NMG^+^ abolished nigericin-mediated NLRP3 activation as measured by IL-1β release (Fig. 7H) and caspase-1 cleavage (Fig. 7I), indicating a requirement for an increase in cytosolic Na^+^. To confirm this, we measured cellular Na^+^ following incubation with nigericin in the presence or absence of extracellular Na^+^. Of note, we could not perform imaging of cytosolic sodium levels as ING-2 is sensitive to acidic pH, which would be induced by nigericin treatment. Strikingly, nigericin triggered an increase in cellular Na^+^ (Fig. 7J), which was completely prevented when extracellular Na^+^ was replaced with NMG^+^ (Fig. 7J). In contrast, substituting Na^+^ for NMG^+^ had no effect on nigericin-mediated K^+^ efflux, demonstrating that the increase in cytosolic sodium does not alter nigericin-mediated cellular K^+^ efflux (Fig. 7K).

Until now, NLRP3 activation by nigericin was considered to be potassium dependent based on the observation that increasing extracellular K^+^ prevented nigericin-driven NLRP3 activation ^38^. Accordingly, we confirmed that increasing extracellular KCl from 5 to 30 mM almost completely inhibited NLRP3 activation by nigericin (Fig. S8E). However, 30 mM extracellular KCl had no effect on the increase in cellular Na^+^ (Fig. S8F), and notably had little effect on cellular K^+^ (Fig. S8G). These experiments demonstrate that, similar to TcdB, increasing extracellular KCl inhibits NLRP3 activation independent of any effect on intracellular ion composition. We also investigated whether the Na^+^-dependent, nigericin-mediated NLRP3 activation involves changes in cell volume. It has been shown previously that increasing the osmolarity of the extra-cellular solution, which we demonstrated to prevent cell swelling, inhibited nigericin-mediated NLRP3 activation ^31^. We assessed the cell volume over time in LPS-primed nigericin-treated hMDM and showed that nigericin triggered an increase in cell volume (Fig. 7L). This was Na^+^-dependent, as the cell volume was reduced when extracellular Na^+^ was substituted with NMG^+^ (Fig. 7L). To determine if the nigericin-dependent cell volume increase was necessary for NLRP3 activation, we incubated hMDM with increasing concentrations of mannitol to boost extracellular osmolarity, which inhibited IL-1β release (Fig. 7M) and caspase-1 cleavage in a concentration-dependent manner (Fig. 7N). It partially reduced the increase in cellular Na^+^ (Fig. S8F), but did not alter the nigericin-dependent decrease in cellular K^+^ (Fig. S8G). Collectively this demonstrates that cytosolic sodium enables nigericin-mediated NLRP3 activation by triggering cell swelling.

## Discussion

Our study establishes that accumulation of cytosolic sodium, triggered by a diverse range of danger associated molecules, is a cellular danger signal that activates the NLRP3 inflammasome. The acute increase in cytosolic sodium activates NLRP3 through at least two distinct pathways: It increases the ion permeability of the endolysosomal membrane, which alters the ionic balance of the cytosol to change cell volume, and it inhibits endosomal trafficking by preventing a decrease in endosomal membrane tension (Fig. S9). Given the low sodium concentration in the cytosol relative to both the endocytic and extracellular compartments, detecting cytosolic sodium accumulation is an effective way to detect permeabilization or ‘leakiness’ of membranes. As sodium can also be removed from the cytosol as it accumulates, it will only activate NLRP3 if the increase is sufficiently large and acute, ensuring minor insults do not lead to inflammasome activation and inflammation. Thus, similar to translocation of endogenous DNA and RNA from the mitochondria, translocation of ions to the cytosol is a danger signal for the cell.

In addition to the initial sodium influx from the endolysosomal compartments, TcdB-mediated NLRP3 activation requires sodium influx from the extracellular space, demonstrating that cytosolic sodium accumulation must cross a threshold in order to activate NLRP3. The TcdB-mediated sodium influx from the extracellular space occurs through macropinocytosis. How macropinocytosis facilitates cytosolic sodium influx is unclear. There are two possibilities: Either by direct import of sodium into the cytoplasm during macropinosome resolution ^37^, or by increasing the plasma membrane localisation of sodium channels or exchangers, which occurs when clathrin-mediated endocytosis is inhibited ^40^. The first explanation seems to be more plausible, as sodium influx mediated by plasma membrane channels would occur in seconds, rather than in minutes as we observe. This indicates that sodium, which is internalized by macropinocytosis and not efficiently removed from the cytoplasm, is interpreted as a danger signal. This is consistent with ouabain-mediated NLRP3 activation, where a failure to remove homeostatic sodium efflux from lumen of macropinosomes enables optimal NLRP3 activation (Fig. S9). This has implications for pathogen detection by NLRP3, as virulence factors that constitutively activate Rac2/3, and so trigger ongoing, uncontrolled macropinocytosis, activate NLRP3 ^41^. This also has repercussions for detection of large clostridial toxins through sodium influx, as all large clostridial toxins inhibit Rac2/3, and subsequently inhibit macropinocytosis once the effector domain reaches the cytosol. As such, NLRP3 activation of large clostridial toxins will depend on how much toxin is internalised, and whether it triggers sufficient sodium translocation to the cytosol prior to inhibition of Rac2/3.

Potassium efflux has long been considered an initiating trigger for NLRP3 activation, as many NLRP3 activators decrease cellular potassium ^38^. However, whether potassium efflux is required for NLRP3 activation, and if so, how it enables NLRP3 activation remains unclear. Here we demonstrate that nigericin, an ionophore that exchanges potassium and hydrogen, activates NLRP3 by triggering a large cellular sodium influx (Fig. S9). As nigericin cannot transport sodium directly into the cell, the increase in cytosolic sodium is likely to be driven by potassium efflux. As potassium is a key regulator of plasma membrane polarization ^42^, potassium efflux could drive sodium influx by activating sodium-permissible ion channels on the plasma membrane through plasma membrane polarization. This is consistent with the requirement for extracellular sodium and the independence of nigericin-mediated NLRP3 activation on macropinocytosis. Candidate channels including either voltage-sensitive sodium channels or transient receptor potential channels, which both open in response to changes in membrane polarization ^43^. Alternatively, sodium hydrogen exchangers in the plasma membrane have also been implicated in cytosolic sodium influx ^40^, and would be activated by the nigericin-dependent acidification of the cytosol. As both sodium channels and exchangers have been the subject of extensive pharmacological research, identification of the sodium influx mechanism and its inhibition may provide scope for the development of new NLRP3 inhibitors.

In comparison to nigericin, the requirement for cellular potassium efflux for activation of NLRP3 by TcdB, ouabain or, indeed, any NLRP3 activator that allows simultaneous sodium influx and potassium efflux, is less clear. We attempted to determine the role of potassium efflux for both TcdB and nigericin by increasing extracellular potassium concentration, which was used in previous studies on NLRP3 ^38^. However, while increasing extracellular potassium to 30 mM robustly inhibits both TcdB- and nigericin-mediated NLRP3 activation, it does so without increasing intracellular potassium. As 30 mM K^+^ is only 20% of the resting intracellular K^+^ concentration, inhibition of NLRP3 by this concentration would imply that almost all K^+^ would have to leave the cell to trigger NLRP3 activation. Furthermore, 30 mM K^+^ does not alter the cytosolic sodium influx, and so inhibits NLRP3 without preventing any of the ionic changes caused by NLRP3 activators. Instead, it is more likely that the increase in extracellular K^+^ inhibits NLRP3 by altering membrane polarization. Increasing extracellular potassium decreases potassium movement across the plasma membrane, which results in a decrease in the plasma membrane polarization. Indeed, increasing the extracellular K^+^ concentration from 5 mM to the 30 mM will decrease the polarization of the plasma membrane by approximately 280% according to the Nerst equation. Therefore, we suggest that inhibition of NLRP3 activation by increased extracellular K^+^ indicates sensitivity towards membrane depolarization rather than potassium efflux.

An important factor for NLRP3 activation is the change of cell volume, and most NLRP3 activators trigger osmotic stress ^44^. Here, we reveal that cytosolic sodium accumulation is a key requirement to induce osmotic stress through cell swelling and activate NLRP3. Importantly, inhibiting cell swelling prevents NLRP3 activation subsequent to any changes in either sodium or potassium. Similarly, hypo-osmotic media-mediated NLRP3 activation is inhibited by preventing change in cell volume even in the presence of decreased cellular potassium ^31,45^. Collectively, this establishes that changes in cellular ionic composition activate NLRP3 in part by their effect on cell volume. How these changes enable NLRP3 activation still need to be determined. Alterations in cell volume will result in remodeling of the plasma membranes ^46^, which may feedback on the Golgi and endosomal compartments, which are suggested as sites of NLRP3 inflammasome assembly. Sodium may also regulate cellular chloride efflux, which also regulates NLRP3 activation ^45,47–49^. Although a 10-20% increase in cell volume caused by TcdB, nigericin, or hypoosmotic media might not be large, small changes in cell volume are known to have a large impact on cellular biology in other contexts. One example is the differentiation of mesenchymal stem cells to osteoclasts, which is regulated by cell-spreading-based volume loss of between 10 and 20% ^50^. Furthermore, cell swelling and subsequent volume changes are regulated primarily by the force of the plasma membrane pushing against the cytoskeleton ^51^, rather than the total area of plasma membrane available. Therefore, a small amount of cell swelling may be sufficient to create the force required to alter cell volume in cells with a highly organized cytoskeleton.

An additional role for cytosolic sodium accumulation in NLRP3 activation occurs through osmotic inhibition of endosomal trafficking. Inhibition of endocytic trafficking, also shown through dispersal of the TGN marker TGN46, has been implicated in NLRP3 activation ^12,13^. However, this has been almost exclusively investigated using ionophores or small molecules that directly disrupt ion gradients across endosomal and TGN membranes and not with physiologically relevant NLRP3 activators, which do not act directly on endosomes or Golgi ^11^. Thus, it was unclear how these activators could perturb either of these compartments. Here, we demonstrate that TcdB-mediated cytosolic sodium accumulation inhibits endocytic trafficking by preventing the osmotic resolution of endosomes and macropinosomes. Sodium accumulation in the cytosol disrupts sodium efflux from the endosomal lumen, preventing water efflux, and the loss of endosomal membrane tension required for onward trafficking ^33^. Importantly, triggering water loss from the lumen of macropinosomes enabled macropinosome resolution even in the presence of increased cytosolic sodium, demonstrating that macropinosome resolution was blocked by osmosis and not the decrease in cytosolic potassium. As many NLRP3 activators, including ATP, pore-forming toxins, and inhibition of the Na^+^/K^+^ ATPase will trigger an increase in cytosolic sodium increase, we suggest it will be the common mechanism to disrupt endocytic trafficking, thereby enabling NLRP3 activation. This finding has also implications beyond NLRP3 activation, as efficient endocytic trafficking is essential for many cellular functions. Thus, our study demonstrates mechanisms through which cytosolic sodium accumulation affects cellular functions, and that a sodium overload can act as a danger signal to inhibit endocytic trafficking and facilitate activation of the NLRP3 inflammasome.

While we have focused on the osmotic effects of sodium on cell volume and endosomal trafficking, it is also possible that sodium may also facilitate NLRP3 activation by having a direct effect on NLRP3 itself. However, as no sodium binding-domains have been identified in any protein so far, and almost all known effects of sodium are through regulation of membrane polarization, organization, or cellular osmolarity ^52,53^, we consider it unlikely that the effects we observe are due to direct action of sodium on NLRP3 and favor the activation by changes in cell volume.

Activation of NLRP3 by cytosolic sodium accumulation provides some explanation for the physiological role for NLRP3 in pathogen detection, which has been controversial^54^. Translocation of pathogen-derived toxins and effectors across the endolysosomal membrane is a highly conserved mechanism used to access the cytosol^55^. Monitoring insertion of these toxins/effectors by detecting the subsequent sodium translocation is advantageous as it enables detection of a wide range of effectors regardless of their other activities. In the case of LCT, NLRP3 is activated by LCT insertion into the endo/lysosome membrane, which is conserved amongst the entire LCT family^4^. Furthermore, the TcdB translocation or delivery domain is conserved across 1104 bacterial proteins expressed by both pathogenic and commensal bacteria proteins, with many proteins also containing toxin-like effector domains^56^. This suggests that the TcdB translocation domain is a common delivery system for bacterial effectors with the potential to disrupt endolysosomal membranes and enable cytosolic sodium influx^57^. A plethora of other pathogenic molecules, including viroporins produced by the influenza virus and SARS-COV-2, as well as bacterial toxins including listeria lysin O and VacA, can potentially enable ion translocation to the cytoplasm. This extends to structurally unrelated toxins, including C3-like toxins that enter the cell by triggering lysosomal swelling and subsequent disruption^58^. Thus, our finding that NLRP3 is a sensor for endolysosomal perturbation by monitoring cytosolic sodium influx provides a framework to understand how NLRP3 detects pathogenic molecules, and should lead to a greater understanding of the role of NLRP3 in pathogen detection in the future.

## Acknowledgments

We thank H. Tatge for purification of the different *C.difficile* toxins. We thank the Microscopy Core Facility and Flow Cytometry Core Facility at the University of Bonn for providing help, service and devices funded by the Deutsche Forschungsgemeinschaft (DFG – German Research Foundation) – Projektnummer 388159768 to the M.C.F. We thank Dr. Andreas Lindner for assistance with analysis of the sodium imaging data. We thank Prof. Stefan Matile and Lea Assies for the EE-flipper dye, Prof. Bernardo Franklin for reagents and the CASY cell analyser, Prof. Veit Hornung for the BLaER1 cells, Prof. Feng Shao for the LFn-PrgI construct, Dr. David Fußböller and Prof. Matthias Geyer for production of the LFn-PrgI. We thank Dr. Nicolas Manel, Dr. Matthieu Piel, Dr. Felix D. Weiss and Dr. Tomasz Prochnicki for scientific discussions. This study was partially funded by the Deutsche Forschungsgemeinschaft (DFG, German Research Foundation) TRR237 (project-ID 369799452), SFB1454 (project-ID 432325352), SFB1403 (project-ID 414786233) and by Germany’s excellence strategy – EXC2151 (project-ID 390873048) to E.L.. We further received funding from the Helmholtz-Gemeinschaft, Zukunftsthema ‘Immunology and Inflammation’ (ZT-0027). In addition, it was partially supported by DFG TRR 374 (project-ID 509149993) and DFG SFB1607 (project-ID 501530074) to J.J. as well as DFG grants GE1017/5-1 and GE1017/5-2 to R.G..

## Author contributions

M.S.J.M. conceived the project. M.S.J.M, D.W., J.J. and E.L. provided scientific input. M.S.J.M., A.A., S.R., A.S., G.H., P.W., and J.S. performed experiments, M.S.J.M., S.R. and G.H. performed data analysis and data interpretation. R.K. and F.D. provide technical support. J.H. performed the electron microscopy. E.L. provided funding for the study. R.G. provided reagents. M.S.J.M and D.W. wrote the manuscript with input from the other authors. M.S.J.M supervised the study and manuscript preparation.

## Supplemental information

Figures S1-S10.

Video S1, LPS treated hMDM, related to Figure 3A-C.

Video S2, LPS and BafA1 treated hMDM, related to Figure 3A-C.

Video S3, LPS and TcdB NXN treated hMDM, related to Figure 3A-C.

Video S4, LPS, TcdB NXN and BafA1 treated hMDM, related to Figure 3A-C.

Video S5, LPS treated hMDM in Na^+^ containing BSS, related to Figure 3G-H.

Video S6, LPS treated hMDM in NMG^+^ containing BSS, related to Figure 3G-H.

Video S7, LPS and TcdB NXN treated hMDM in Na^+^ containing BSS, related to Figure 3G-H.

Video S8, LPS and TcdB NXN treated hMDM in NMG^+^ containing BSS, related to Figure 3G-H.

Video S9, LPS treated hMDM, related to Figure 4D-G.

Video S10, LPS and TcdB NXN treated hMDM, related to Figure 4D-G.

Video S11, LPS and TcdB NXN treated hMDM preincubated with EIPA, related to Figure 4D-G.

Video S12, LPS and TcdB NXN treated hMDM preincubated with LY294002, related to Figure 4D-G.

Video S13, LPS and TcdB NXN treated hMDM preincubated with NPS2143, related to Figure 4D-G.

Video S14, LPS treated hMDM assessed for macropinosome maturation, related to Figure 6F, Figure S7D.

Video S15, LPS and TcdB NXN treated hMDM assessed for macropinosome maturation, related to Figure 6F, Figure S7D.

Video S16, LPS treated hMDM assessed for macropinosome maturation, related to Figure S7E.

Video S17, LPS and TcdB NXN treated hMDM assessed for macropinosome maturation, related to Figure S7E. Timing of the hyperosmotic pulse is anno- tated in the Video.

