## Supplementary_figures for "Cytosolic sodium accumulation is a common danger signal triggering endocytic dysfunction and NLRP3 inflammasome activation"

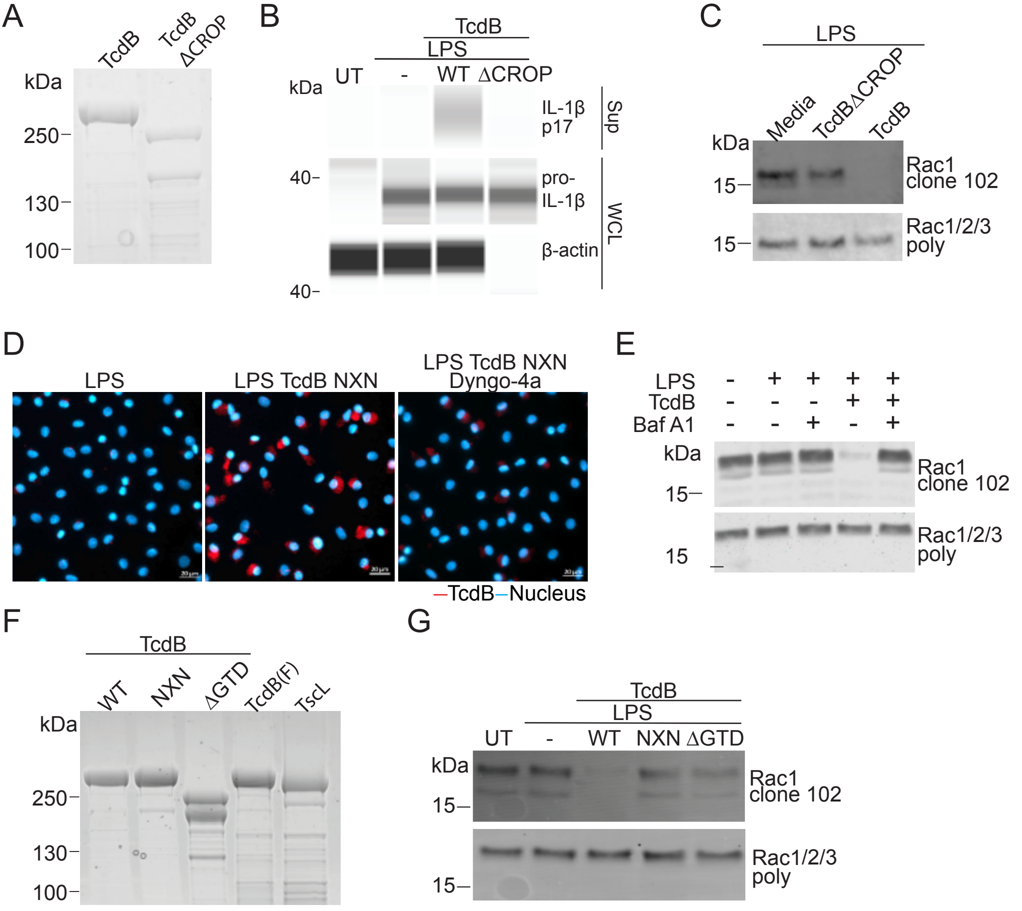


#### Supplementary figure 1

**A)** Coomassie stained SDS-PAGE gel characterizing TcdB and the TcdB ΔCROP mutant. LPS-primed hMDM (10 ng/ml, 3 h) incubated with TcdB or TcdB ΔCROP (20 ng/ml, 2h). IL-1β assessed from sup or WCL by SWA **(B)** or TcdB-modified Rac (Rac antibody clone 102) or total Rac (Rac1/2/3 poly) assessed by immunoblot from WCL **(C)**. **D)** Fluorescence imaging of hMDM treated as in Fig. 1D and stained for TcdB (A647) and nuclei (DAPI). **E)** Rac2/3 modification assessed by immunoblot from WCL from hMDM treated as in Fig 1E. **F)** Coomassie stained SDS-PAGE gel characterizing TcdB, TcdB mutants and TscL. **G)** Rac2/3 modification assessed by immunoblot from WCL from hMDM treated as in Fig 1F. A) and F) n=1, B-E) and G) n=2.

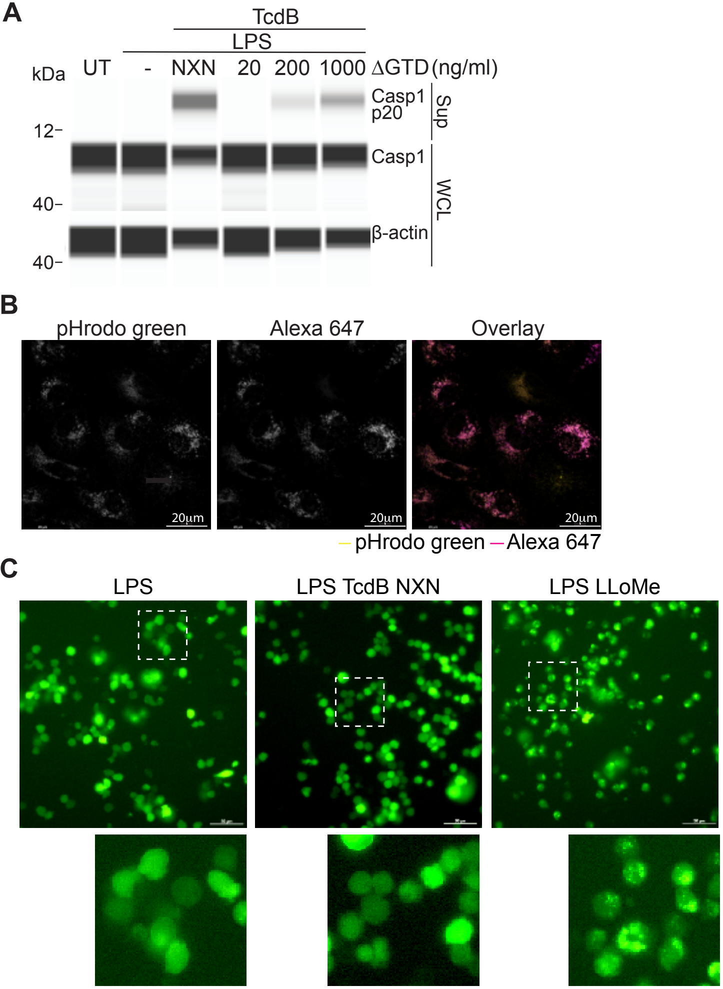


**Supplementary figure 2**

**A)** LPS-primed hMDM and incubated with TcdB NXN (20 ng/ml, 2 h), or with increasing concentrations of TcdB ΔGTD (2 h). Caspase-1 was assessed from sup or WCL by SWA, representative of n = 2 **B)** Fluorescent confocal images of hMDM loaded with pHrodo green- or Alexafluor 647-coupled 10 kDa dextrans. Representative of 2 independent experiments **C)** Fluorescent images of trans-differentiated BLaER1 cells expressing a Gal-8-mCitrine fusion protein were primed with LPS (100 ng/ml, 3h) and treated with either TcdB NXN (20 ng/ml, 45 min) or LLoMe (0.5 mM, 30 min). Representative of n= 3.

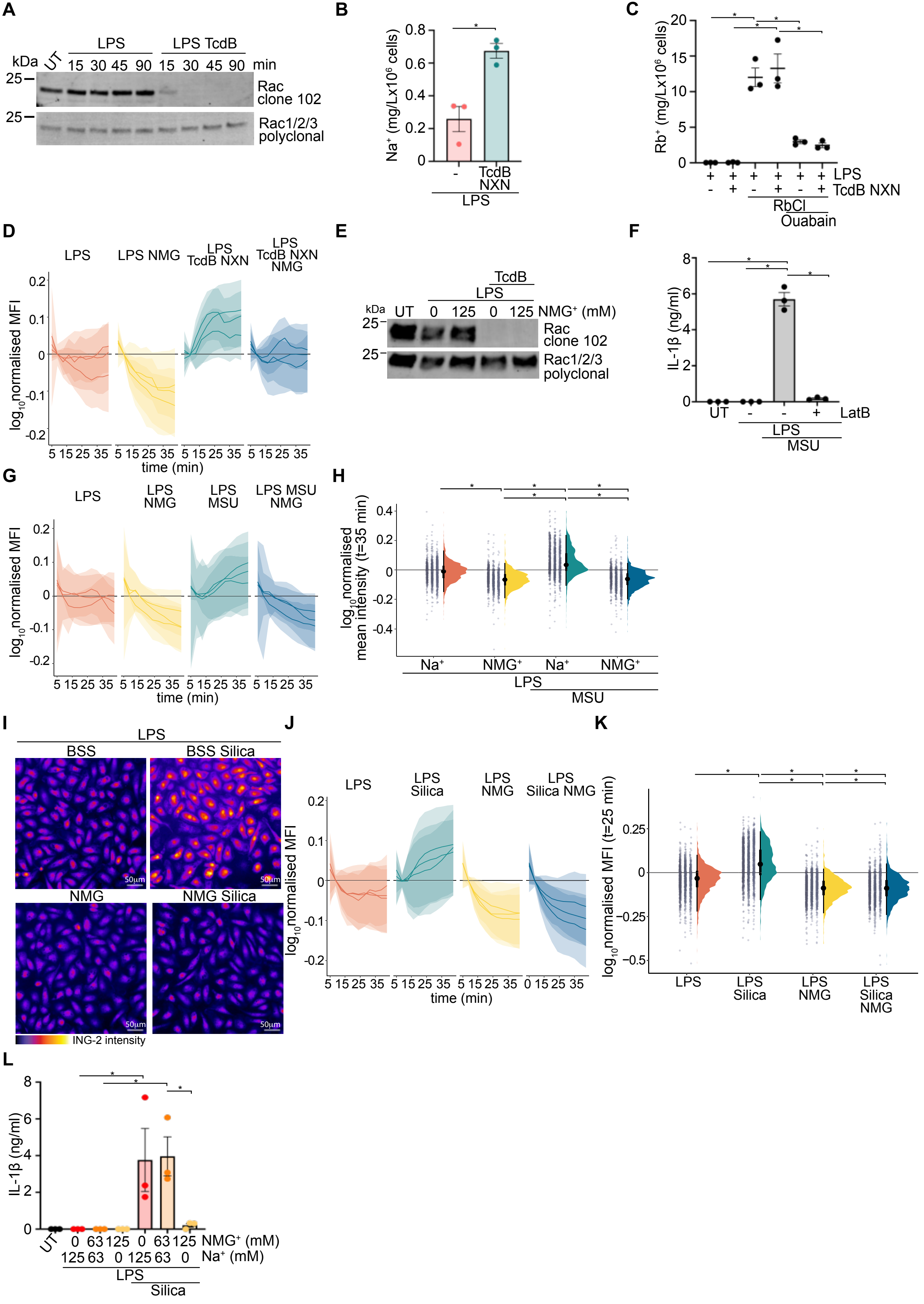
A
LPS
LPS TcdB
kDa
UT
15 30 45 90 15 30 45 90 min
25
Rac clone 102
25
Rac1/2/3 polyclonal
B
Na⁺ (mg/Lx10⁶ cells)
-
TcdB NXN
LPS
C
Rb⁺ (mg/Lx10⁶ cells)
LPS
TcdB NXN
RbCl
Ouabain
D
LPS
LPS NMG
LPS TcdB NXN
LPS TcdB NXN NMG
log10normalised MFI
time (min)
E
TcdB
LPS
kDa
UT
0 125 0 125 NMG⁺ (mM)
Rac clone 102
Rac1/2/3 polyclonal
F
IL-1β (ng/ml)
UT
LatB
LPS
MSU
G
LPS
LPS NMG
LPS MSU
LPS MSU NMG
log10normalised MFI
time (min)
H
log10normalised mean intensity (t=35 min)
Na⁺
NMG⁺
Na⁺
NMG⁺
LPS
MSU
I
LPS
BSS
BSS Silica
NMG
NMG Silica
50µm
ING-2 intensity
J
LPS
LPS Silica
LPS NMG
LPS Silica NMG
log10normalised MFI
time (min)
K
log10normalised MFI (t=25 min)
LPS
LPS Silica
LPS NMG
LPS Silica NMG
L
IL-1β (ng/ml)
UT
0 63 125 0 63 125 NMG⁺ (mM)
125 63 0 125 63 0 Na⁺ (mM)
LPS
Silica

### Figure S3

**Supplementary figure 3**

**A)** Immunoblot of Rac2/3 from lysate of LPS-primed hMDM incubated with TcdB, representative of n=3. **B)** Cellular $Na^+$ from LPS-primed hMDM +/- TcdB NXN (20 ng/ml, 45 min). **C)** $Rb^+$ uptake in LPS-primed hMDM +/- TcdB NXN +/- ouabain (2.5 mM) for 1h. **D)** Temporal changes of ING-2 intensity normalised to t = 0 min, per cell for LPS-primed hMDM treated as in Fig. 3G. Mean (solid line) ± S.D. (shaded area) is shown for each experimental replicate. **E)** Immunoblot of Rac2/3 from LPS-primed hMDM +/- TcdB in $Na^+$ or $NMG^+$ containing BSS, representative of n=2. **F)** LPS-primed hMDM pre-incubated with Latrunculin B (LatB) (2 μM, 30 min), then incubated with MSU crystals (100 μg/ml) 2h. IL-1β measured from sup. LPS-primed hMDM treated with MSU crystals (100 μg/ml) in $Na^+$ or $NMG^+$ containing BSS and the **G)** Temporal changes of ING-2 intensity normalised to t = 0 min, per cell. Mean (solid line) ± S.D. (shaded area) is shown for each experimental replicate. **H)** Normalized ING-2 intensities at t = 35 min from G). Grey dot represents one cell, mean ± S.D. are shown. **I)** Fluorescent images of LPS-primed, ING-2 loaded hMDM stimulated with Silica (100 μg/ml) on $Na^+$ or $NMG^+$ BSS. **J)** Temporal changes of ING-2 intensity normalised to t = 0 min, per cell. Mean (solid line) ± S.D. (shaded area) is shown for each experimental replicate. **K)** Normalized ING-2 intensities at t = 35 min from G). Grey dot represents one cell, mean ± S.D. are shown. **L)** IL-1β measured from sup from LPS-primed hMDM stimulated with Silica (100 μg/ml) on $Na^+$ or $NMG^+$ BSS. Mean ± SEM, n = 3, each point represents one donor. hMDM were pre-incubated with VX-765 in B)-D) and G)-K) prior to addition of the inflammasome stimulus. Statistical analysis performed using one-way ANOVA, *$p<0.05$, only significant differences annotated.

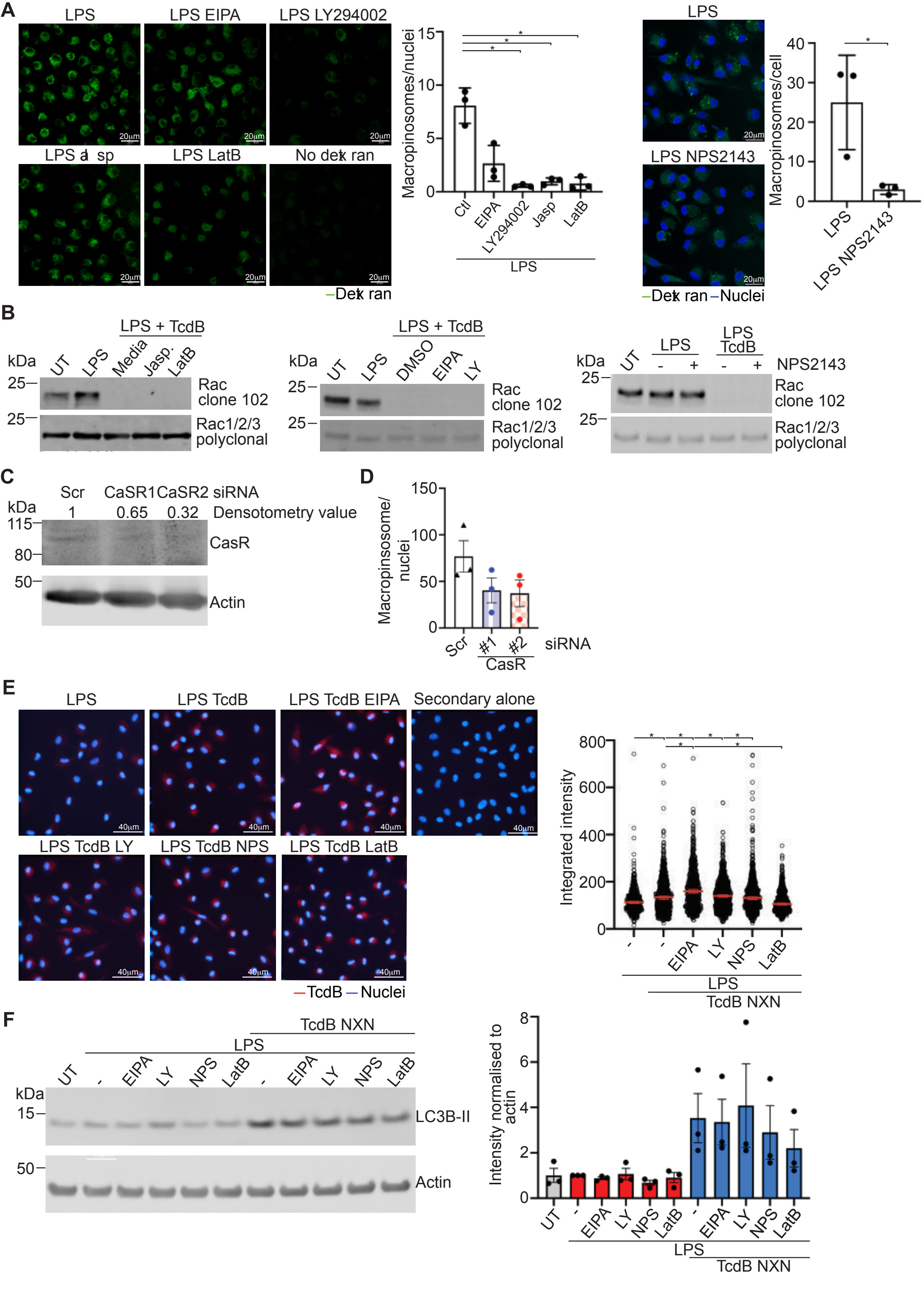
A
LPS
LPS EIPA
LPS LY294002
LPS Jasp
LPS LatB
No dextran
20μm
Dextran
Macropinosomes/nuclei
Ctl
EIPA
LY294002
Jasp
LatB
LPS
LPS
LPS NPS2143
Dextran
Nuclei
Macropinosomes/cell
B
LPS + TcdB
kDa
UT
LPS
Media
Jasp.
LatB
Rac clone 102
Rac1/2/3 polyclonal
DMSO
EIPA
LY
NPS2143
C
Scr
CaSR1
CaSR2 siRNA
Densotometry value
1
0.65
0.32
CasR
Actin
D
Macropinosome/ nuclei
siRNA
CasR
E
LPS TcdB
LPS TcdB EIPA
Secondary alone
LPS TcdB LY
LPS TcdB NPS
LPS TcdB LatB
40μm
TcdB
Nuclei
Integrated intensity
TcdB NXN
F
LC3B-II
Actin
Intensity normalised to actin
NPS

**Supplementary figure 4**

Fluorescence confocal images and analysis of LPS-primed hMDM incubated with EIPA (20 μM), LY294002 (10 μM), Jasplakinolide (0.5 μM), Latrunculin B (2 μM) or NPS2143 (5 μM) for 30 min, then incubated with OG-labeled 70kDa dextran for 15 min (25 μg/ml). The mean number of macropinosomes was quantitated for each condition (n>100 cells per condition). Mean ± SEM, n = 3, each point represents one donor. **B)** Rac2/3 modification assessed from WCL by immunoblot from hMDM treated with inhibitors as in Fig 4B, then incubated with TcdB (20 ng/ml) for 45 min. Representative of n = 3. **C)** CaSR expression assessed from WCL by immunoblot from CaSR or scr siRNA treated hMDM. Representative of n = 2. **D)** Quantitation of macropinosomes in hMDM transfected with CaSR targeting or Scr targeting siRNA. Mean ± SEM, n = 3, 30-50 cells analysed per donor. Each point represents data from one donor. **E)** Fluorescence imaging and analysis of TcdB uptake in LPS-primed hMDM incubated with inhibitors as in Fig. 4B and incubated with TcdB NXN for 45 min in the presence of CP-456,773. Mean ± 95% CI shown, representative of n = 2. **F)** LC3B-II levels and quantitation in hMDM treated as in Fig. 4B, mean ± SEM, n = 3, each point represents one donor. Statistical analysis performed using one-way ANOVA, *p<0.05, only significant differences annotated.

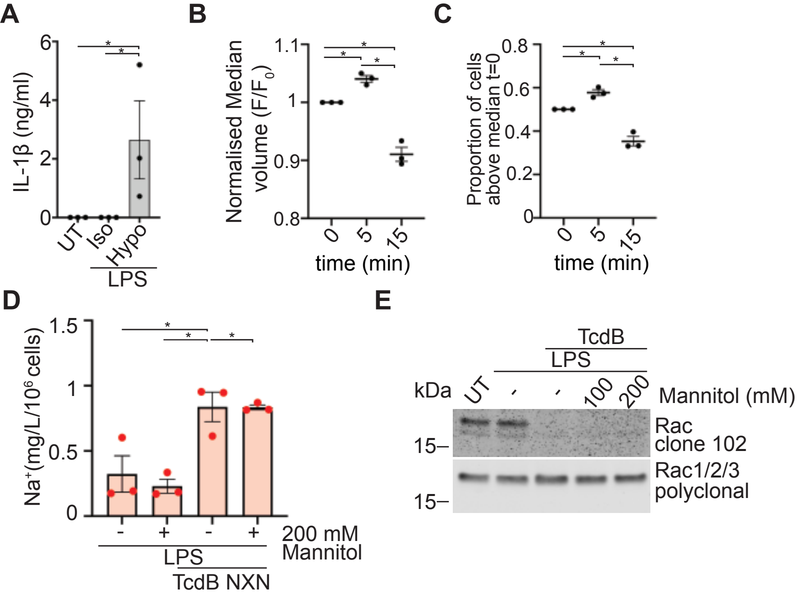


**Supplementary figure 5**

**A)** LPS-primed hMDM incubated with either isotonic or hypotonic BSS for 2h, IL-1β measured from sup. **B)** Normalised cell volume or **C)** the proportion of hMDM above the median from t=0 from LPS-primed hMDM incubated in hypoosmotic media for 0 min, 5 min, or 15 min. Mean ± SEM, n = 3, each point represents one donor. **D)** Cellular $Na^+$ from LPS-primed hMDM +/- TcdB NXN (20 ng/ml, 45 min) in media +/- 200 mM mannitol. Performed in the presence of VX-765. Mean ± SEM, n = 3, each point represents one donor. **E)** Immunoblot of Rac2/3 from lysate of LPS-primed hMDM incubated with TcdB with 100 mM or 200 mM mannitol. Representative of n = 2. Statistical analysis performed with one-way ANOVA, $^*p<0.05$, only significant differences annotated.

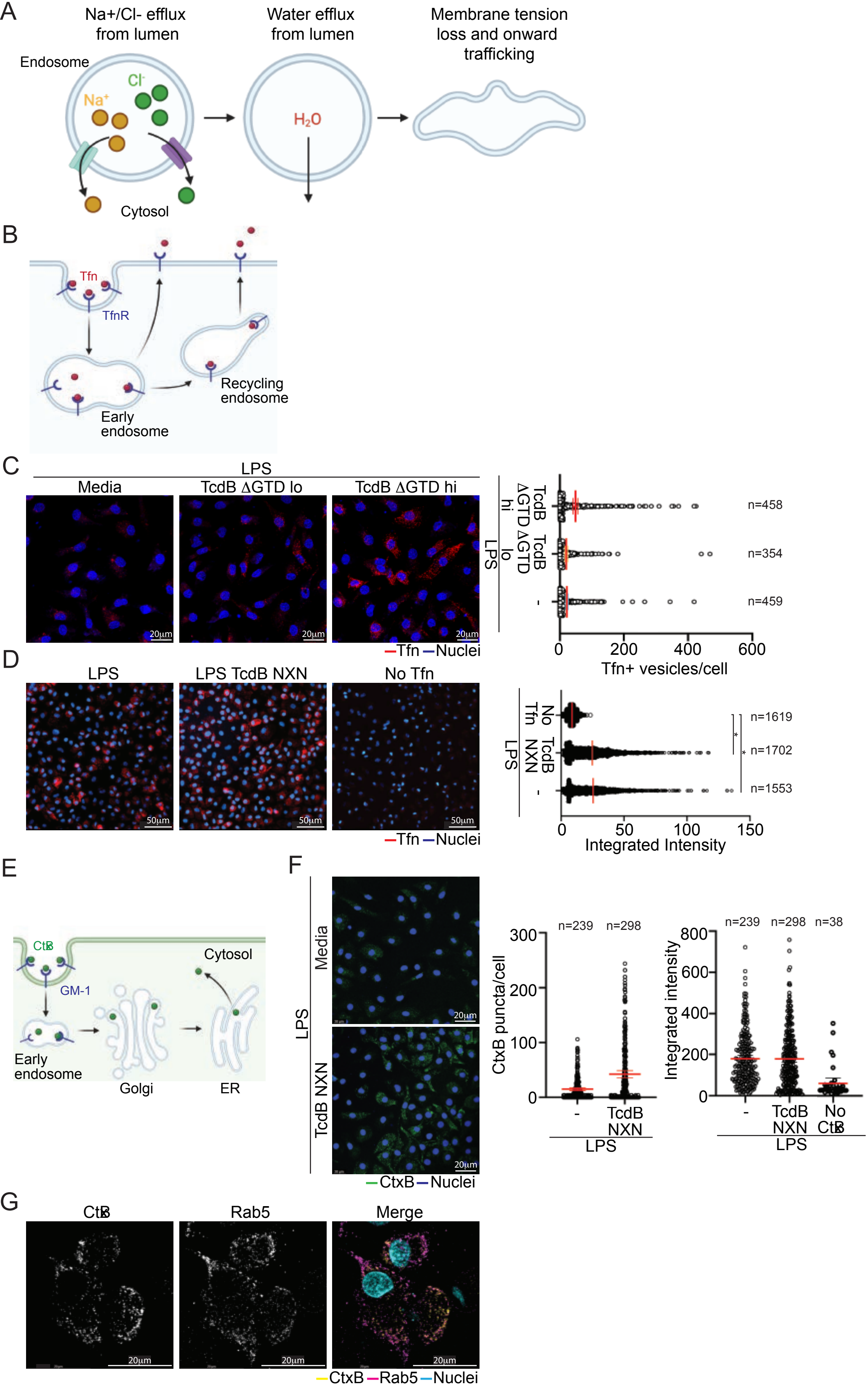
A
Na+/Cl- efflux from lumen
Water efflux from lumen
Membrane tension loss and onward trafficking
Endosome
Cl-
Na+
H2O
Cytosol
B
Tfn
TfnR
Recycling endosome
Early endosome
C
LPS
Media
TcdB ΔGTD lo
TcdB ΔGTD hi
20µm
Tfn Nuclei
n=458
n=354
n=459
Tfn+ vesicles/cell
D
LPS
LPS TcdB NXN
No Tfn
50µm
Tfn Nuclei
n=1619
n=1702
n=1553
Integrated Intensity
E
CtxB
GM-1
Cytosol
Early endosome
Golgi
ER
F
Media
TcdB NXN
CtxB Nuclei
n=239
n=298
n=38
CtxB puncta/cell
Integrated intensity
No CtxB
G
CtxB
Rab5
Merge
CtxB Rab5 Nuclei

**Supplementary figure 6**

**A)** A schematic of the osmotic resolutions of endosomes. **B)** A schematic of the Tfn trafficking pathway. **C)** Fluorescence confocal images and analysis of Tfn-A647 pulse-chase (chase period 30 min) in LPS-primed hMDM incubated with TcdBΔGTD lo (20 ng/ml) or TcdBΔGTD hi (1 μg/ml). Mean ± 95% CI, each point represents one cell, data pooled from three independent experiments. **D)** Fluorescence imaging and analysis of Tfn binding for LPS-primed hMDM. Mean ± 95% CI, each point represents one cell, representative of two independent experiments. **E)** Schematic of the CtxB trafficking pathway. **F)** Fluorescence confocal images and analysis of CtxB-A555 pulse-chase (pulse 10 min, chase period 30 min) in LPS-primed hMDM incubated with TcdB NXN preincubated with CP-456,773. Mean ± 95% CI, each point represents one cell, data pooled from three independent experiments. **G)** Fluorescence confocal images of CtxB-A555 and Rab5 in LPS-primed, TcdB NXN treated hMDM. Representative of approx. 50 cells per donor for 3 donors. C), F) and G) were performed in the presence of CP-456,773. Statistical analysis performed using one-way ANOVA, *$p<0.05$, only significant differences annotated.

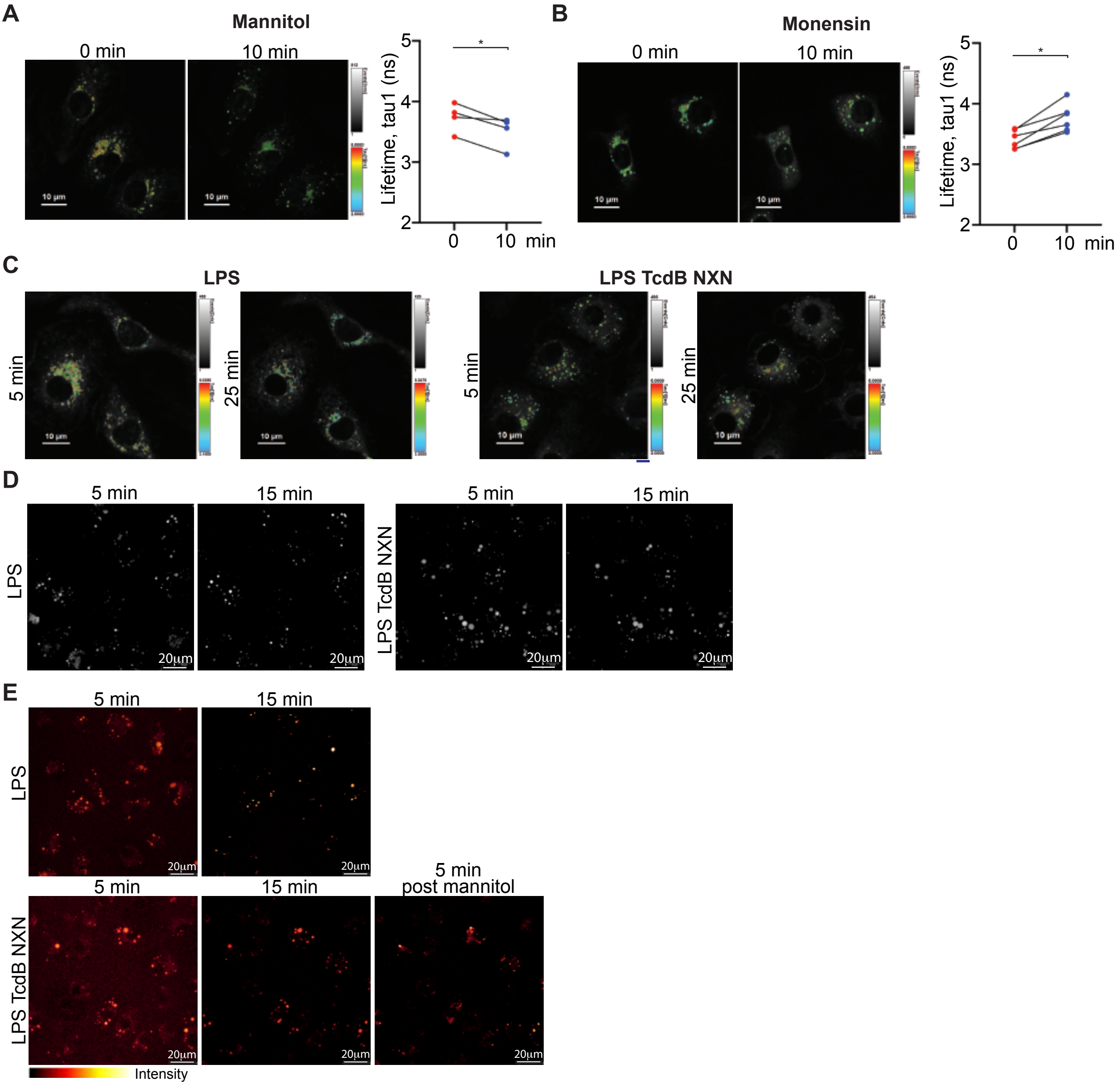


#### Supplementary figure 7

FLIM images and analysis for EE-flipper (2 μM, 10 min) loaded, LPS-primed hMDM treated with **A)** mannitol (0.25 M), **B)** monensin (2 μM) for 10 min or TcdB for 25 min. Each pair represents one experimental repeat. **C)** FLIM images for EE-flipper (2 μM, 10 min) loaded, LPS-primed hMDM treated as in Fig. 5D. **D)** Greyscale confocal images of the images shown in Fig 5F. **E)** LPS-primed hMDM +/- TcdB NXN treated as in Fig. 5F were trreated with 0.2 M mannitol 15 min post-dextran pulse, then imaged for a further 5 min. Representative of 3 independent experiments. C), D) and E) were performed in the presence of CP-456,773. Statistical analysis performed using a paired Student's t-test. *p<0.05, only significant differences annotated.

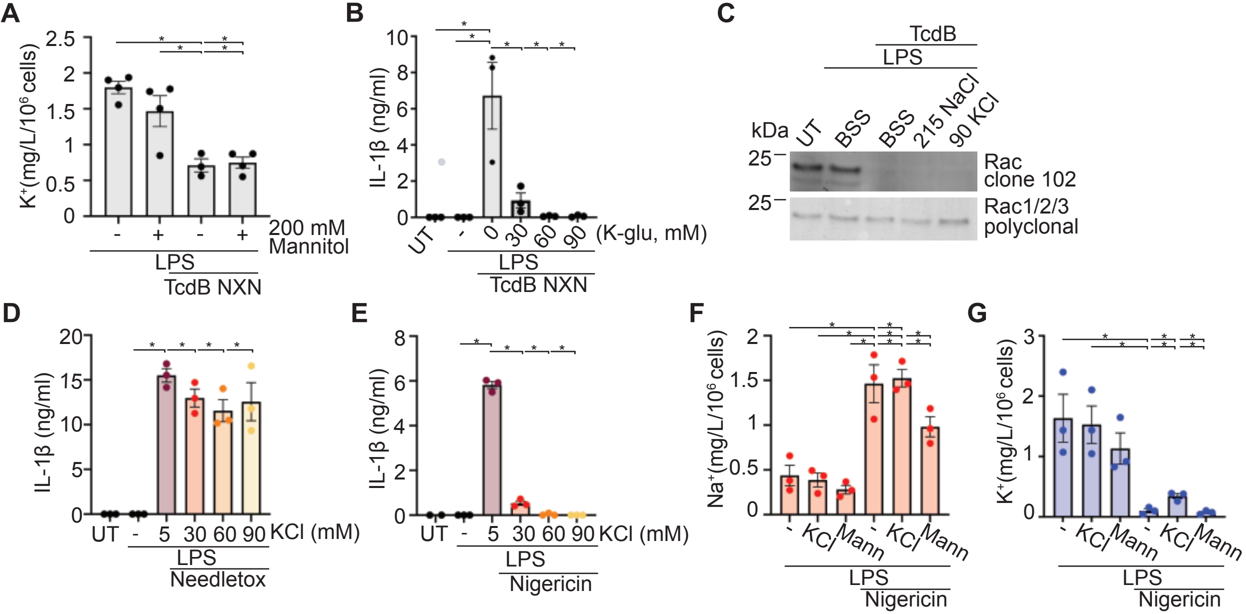


**Supplementary figure 8**

**A)** Cellular $K^+$ from LPS-primed hMDM +/- TcdB NXN (20 ng/ml, 45 min) in media containing 200 mM mannitol. **B)** LPS-primed hMDM incubated with TcdB NXN (20 ng/ml, 2 h) in media containing increasing concentrations of K-gluconate. IL-1β was measured from sup. **C)** Rac2/3 modification measured from WCL by immunoblot from LPS-primed hMDM incubated with TcdB with either 5 mM or 30 mM extracellular KCl, representative of n = 2. **D)** LPS-primed hMDM incubated with needletox in media containing increasing concentrations of KCl. IL-1β was measured from sup. **E)** LPS-primed hMDM incubated with nigericin (8 μM, 2 h) in media containing increasing concentrations of KCl. IL-1β was measured from sup. Cellular $Na^+$ **F)** or $K^+$ **G)** from LPS-primed hMDM +/- nigericin (8 μM, 45 min) in media containing 5 mM or 30 mM KCl or 200 mM mannitol. Representative of n = 2. Mean ± SEM, n = 3, each point represents one donor. A), F) and G) were performed in the presence of VX-765. Statistical analysis performed with one-way ANOVA, *$p<0.05$, only significant differences annotated.

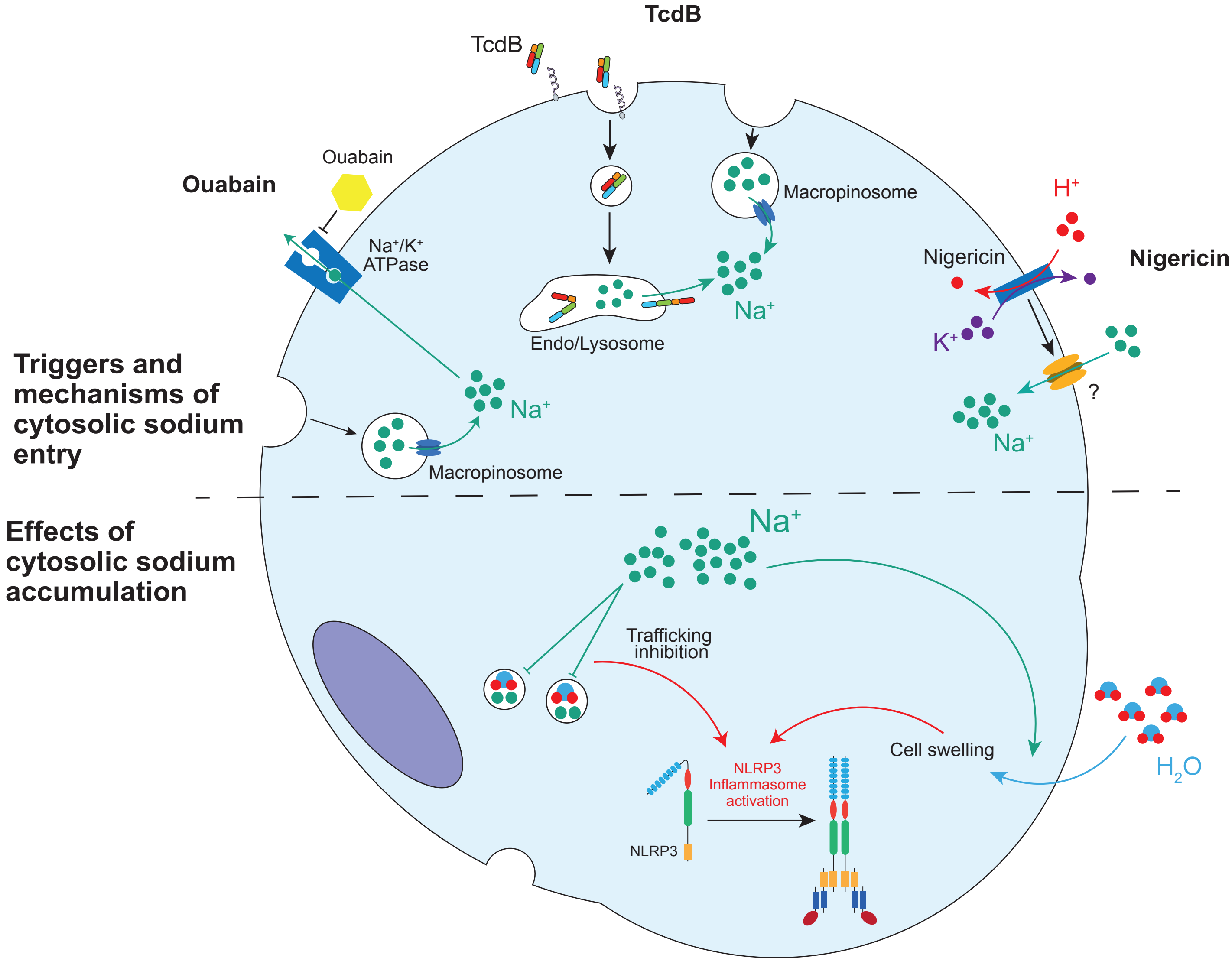


**Supplementary figure 9**

Model of cytosolic sodium influx-mediated NLRP3 activation.

**Supplementary figure 10**

Single color, secondary antibody alone and unloaded controls for **A)** Fig. 2C **B)** Fig. 3A **C)** Fig. 6A, S6F **D)** Fig. 6C **E)** Fig. 6B **F)** Fig. S6G **G)** Fig. 6F **H)** Unstained control for all EE-flipper experiments.

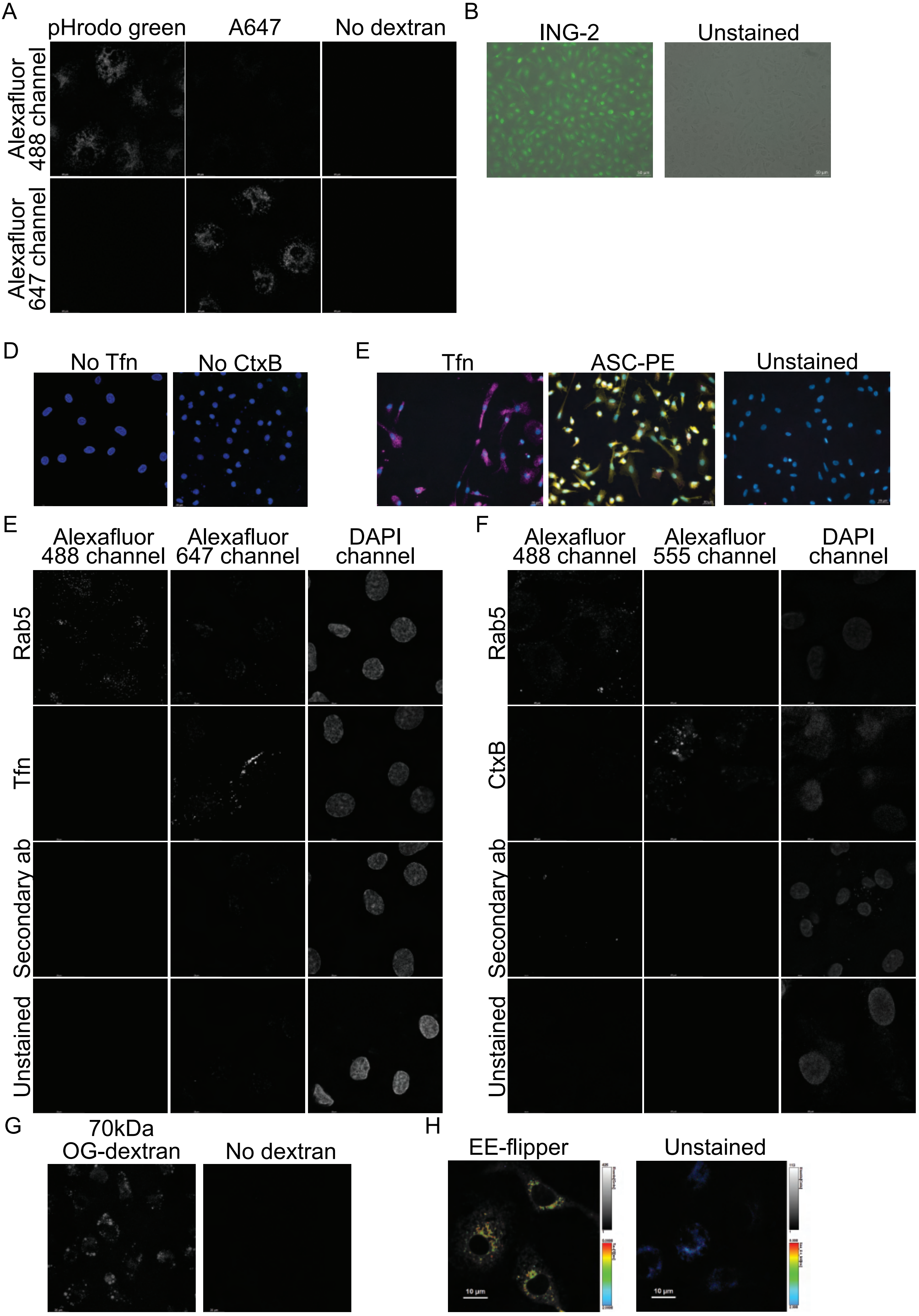
A
pHrodo green
A647
No dextran
Alexafluor 488 channel
Alexafluor 647 channel
B
ING-2
Unstained
D
No Tfn
No CtxB
E
Tfn
ASC-PE
Unstained
E
Alexafluor 488 channel
Alexafluor 647 channel
DAPI channel
Rab5
Tfn
Secondary ab
Unstained
F
Alexafluor 488 channel
Alexafluor 555 channel
DAPI channel
Rab5
CtxB
Secondary ab
Unstained
G
70kDa OG-dextran
No dextran
H
EE-flipper
Unstained
10 µm
10 µm

### Figure S10

**Captions for Movies S1 to S17**

Movies S1-S4, hMDM were from hMDM treated as in Fig. 2A, movies S5-S8 were from hMDM treated as in Fig. S3D, movies S9-S13 were from hMDM treated as in Fig. 3D. Movies S14 and S15 of macropinosome maturation as shown in Fig. 5E, movies S16 and S17 of macropinosomes in hMDM subject to hyperosmotic pulse as in Fig. S7B. All movies were performed in the presence of CP-456,773 (2.5 μM) to ensure that any effects observed were proximal to NLRP3 activation.

**Movie S1**

LPS treated hMDM.

**Movie S2**

LPS and BafA1 treated hMDM.

**Movie S3**

LPS and TcdB NXN treated hMDM.

**Movie S4**

LPS, TcdB NXN and BafA1 treated hMDM.

**Movie S5**

LPS treated hMDM in $Na^+$ containing BSS.

**Movie S6**

LPS treated hMDM in $NMG^+$ containing BSS.

**Movie S7**

LPS and TcdB NXN treated hMDM in $Na^+$ containing BSS.

**Movie S8**

LPS and TcdB NXN treated hMDM in $NMG^+$ containing BSS.

**Movie S9**

LPS treated hMDM.

**Movie S10**

LPS and TcdB NXN treated hMDM.

**Movie S11**

LPS and TcdB NXN treated hMDM preincubated with EIPA.

**Movie S12**

LPS and TcdB NXN treated hMDM preincubated with LY294002.

**Movie S13**

LPS and TcdB NXN treated hMDM preincubated with NPS2143.

**Movie S14**

LPS treated hMDM assessed for macropinosome maturation as in Fig. 5E.

**Movie S15**

LPS and TcdB NXN treated hMDM assessed for macropinosome maturation as in Fig. 5E.

**Movie S16**

LPS treated hMDM assessed for macropinosome maturation as in Fig. S7B.

**Movie S17**

LPS and TcdB NXN treated hMDM assessed for macropinosome maturation as in Fig. S7B. Timing of the hyperosmotic pulse is annotated in the movie.
